# Population and Evolutionary Genetics Subfamily-specific functionalization of diversified immune receptors in wild barley

**DOI:** 10.1101/352278

**Authors:** Takaki Maekawa, Barbara Kracher, Isabel M. L. Saur, Makoto Yoshikawa-Maekawa, Ronny Kellner, Artem Pankin, Maria von Korff, Paul Schulze-Lefert

## Abstract

Gene-for-gene immunity between plants and host-adapted pathogens is often linked to population-level diversification of immune receptors encoded by disease resistance (*R*) genes. The complex barley (*Hordeum vulgare* L.) *R* gene locus *Mildew Locus A* (*Mla*) provides isolate-specific resistance against the powdery mildew fungus *Blumeria graminis* f. sp. *hordei* (*Bgh*) and has been introgressed into modern barley cultivars from diverse germplasms, including the wild relative *H. spontaneum*. Known *Mla* disease resistance specificities to *Bgh* appear to encode allelic variants of the R Gene Homolog 1 (RGH1) family of nucleotide-binding domain and leucine-rich repeat (NLR) proteins. To gain insights into *Mla* diversity in wild barley populations, we here sequenced and assembled the transcriptomes of 50 accessions of *H. spontaneum* representing nine populations distributed throughout the Fertile Crescent. The assembled *Mla* transcripts exhibited rich sequence diversity, which is linked neither to geographic origin nor population structure. *Mla* transcripts in the tested *H. spontaneum* accessions could be grouped into two similar-sized subfamilies based on two major N-terminal coiled-coil signaling domains that are both capable of eliciting cell death. The presence of positively selected sites, located mainly in the C-terminal leucine-rich repeats of both MLA subfamilies, together with the fact that both coiled-coil signaling domains mediate cell death, implies that the two subfamilies are actively maintained in the host population. Unexpectedly, known MLA receptor variants that confer *Bgh* resistance belong exclusively to one subfamily. Thus, signaling domain divergence, potentially to distinct pathogen populations, is an evolutionary signature of functional diversification of an immune receptor.

## Article Summary

Powdery mildew poses a significant threat to barley production worldwide, and *Mildew Locus A* (*Mla*) gene variants have been introgressed into modern cultivars from wild barleys to provide disease resistance. We found that *Mla* genes in wild barley accessions are grouped into two subfamilies with all known variants effective against powdery mildew disease belonging exclusively to one subfamily, suggesting that the two subfamilies have evolved to combat distinct pathogens. Furthermore, divergence of the signalling domain but not the pathogen-recognition domain of MLA receptors defines subfamily grouping, emphasizing the importance of choice of signalling domain haplotype for effective disease control.

## Introduction

Adaptation to pathogens is linked to a range of evolutionary processes that result in genetic variation and affect disease resistance traits in a host population. In the co-evolutionary “arms race” model, recurrent allele fixation in a host population is predicted to reduce genetic diversity, whereas in the “trench warfare” model co-existence of functional and nonfunctional alleles is possible when a fitness penalty is associated with a functional allele in the absence of pathogens (Stahl, et al. 1999; Tian, et al. 2003). In plants, “gene-for-gene” resistance (Flor 1955) is frequently found in interactions between hosts and host-adapted pathogens and is often associated with population-level diversification of immune receptors encoded by disease resistance (*R*) genes (Maekawa, Kufer, et al. 2011). The products of these *R* genes recognize matching pathogen effectors, designated avirulence (AVR) effectors, and plants that lack matching *R* genes are susceptible to effector-mediated pathogen virulence (Jones and Dangl 2006). A single *R* gene can encode functionally diversified resistance alleles among individuals of the host population (Maekawa, et al. 2011). While such balancing selection at a given *R* locus can potentially be due to co-evolutionary diversification of a single *R* and *AVR* gene pair, in some cases it is rather explained by the recognition of multiple non-homologous effectors derived from different pathogens (Karasov, et al. 2014; Anderson, et al. 2016; Lu, et al. 2016).

Individual *R* genes are often members of larger gene families, organized in complex loci of paralogous genes, and can evolve through tandem and segmental gene duplications, recombination, unequal crossing-over, and point mutations (Jacob, et al. 2013). Most known *R* genes encode intracellular nucleotide-binding domain and leucine-rich repeat proteins (NLRs). Plant NLRs belong to a subclass of the STAND (signal transduction ATPases with numerous domains) superfamily of proteins (Maekawa et al., 2011), which possess variable N-terminal domains, a central conserved NB-ARC [Nucleotide-Binding, shared by *A*poptotic protease activating factor 1 (Apaf-1), certain *R*-proteins, and Cell death protein 4 (CED-4)] domain, and C-terminal leucine-rich repeat region (LRR) with varying repeat number. Incorporation of either a TOLL/interleukin 1 receptor (TIR)-like domain or a coiled-coil (CC) domain at the N-terminus defines two major classes of NLRs, designated TNLs and CNLs, respectively. Extensive cross-plant species database searches have revealed that approximately 10% of plant NLRs additionally contain highly variable integrated domains (Ellis 2016). Effector proteins can be recognized directly by NLRs inside plant cells, in what are essentially receptor-ligand interactions (Dodds, et al. 2006; Ortiz, et al. 2017), or, indirectly, through modifications of host proteins that are associated with the NLR receptor (Mackey, et al. 2002; Axtell and Staskawicz 2003; Dodds, et al. 2006; Gutierrez, et al. 2010; Ntoukakis, et al. 2013; Ortiz, et al. 2017). During direct non-self perception, the C-terminal LRR or integrated domains are known to function as major determinants of recognition specificity for AVR effectors (Dodds, et al. 2006; Cesari, et al. 2013; Maqbool, et al. 2015; Zhu, et al. 2017). Collectively, population-level *R* gene diversification and the diversity of NLR-mediated non-self recognition mechanisms maximize the ability of plants to cope with rapidly evolving pathogen effectors.

Powdery mildews are fungal plant pathogens that are ubiquitous in temperate regions of the world and infect nearly 10,000 species of angiosperms (Glawe 2008). Given that the powdery mildew disease caused by *Blumeria graminis* f. sp. *hordei* (*Bgh*) poses a significant threat to barley production (*Hordeum vulgare* L.), powdery mildew *R* loci, present in germplasm collections of barley relatives including wild barley (*H*. *spontaneum* L.), have been extensively investigated (Jahoor and Fischbeck 1987; Jørgensen 1994). These efforts have revealed numerous powdery mildew *R* loci and a subset was subsequently introgressed into modern barley cultivars to genetically control the disease (Jørgensen 1994). Among these, the Mildew resistance Locus A (*Mla)* is characterized by an exceptional functional diversification; each *Mla* locus in a given accession confers disease resistance to a distinctive set of *Bgh* test isolates (races), designated *Mla* resistance specificity (Jørgensen 1994; (Seeholzer, et al. 2010). More than 30 *Bgh* isolate (race)-specific resistance specificities map close or at the *Mla* locus (Jørgensen 1994). A 265-kb contiguous DNA sequence spanning the *Mla* locus in the barley reference cultivar ‘Morex’ consists of a cluster of CNL-encoding genes belonging to three distinctive families, which are designated *R* Gene Homolog (RGH) 1, RGH2 and RGH3 (Wei, et al. 2002). To date, 28 naturally diversified RGH1 sequences have been molecularly characterized and the majority of these are capable of conferring isolate-specific immunity to *Bgh* (Seeholzer, et al. 2010). These sequence variants appear to represent *Rgh1* alleles at the *Mla* locus as evidenced by the presence of an [AT]_n_ microsatellite in the third intron (Shen, et al. 2003). However, the presence of the microsatellite has been validated for the genomic *Rgh1* sequences of only six *Mla* resistance specificities to *Bgh*. Cultivar ‘Morex’ carries a truncated and non-functional *Rgh1* allele, designated *Rgh1bcd* and few cultivars appear to harbor more than one functional *Rgh1* copy (Wei, et al. 2002; Seeholzer, et al. 2010). Our recent work demonstrated that sequence-related MLA receptor variants recognize sequence-unrelated *Bgh* effectors via direct interaction (Lu, et al. 2016). In addition, the wheat *Mla* orthologs *Sr33* (*Stem rust* resistance 33) (Periyannan, et al. 2013) and *Sr50* (*Stem rust* resistance 50) (Mago, et al. 2015), introgressed from *Aegilops tauschii* and *Secale cereale,* respectively, confer resistance to the stem rust pathogen *Puccinia graminis* f. sp. *tritici* (*Pgt*) isolate Ug99, a pathogen that poses a major threat to global wheat production. These findings suggest that the last common ancestor of these cereals harbored an ancestral *Mla* gene and imply that *Mla* diversified to detect multiple non-homologous effectors from at least two unrelated fungal pathogens, the Ascomycete *Bgh* and Basidiomycete *Pgt*.

Overexpression of the MLA, Sr33, and Sr50 N-terminal CC domains is sufficient to initiate immune signaling similar to that mediated by the corresponding full-length receptors (i.e. activation of host cell death and transcriptional reprograming for immune responses; (Maekawa, Cheng, et al. 2011; Casey, et al. 2016; Cesari, et al. 2016; Jacob, et al. 2018). Functional analysis of MLA chimeras suggests that the LRR determines AVR recognition specificities in *Bgh* resistance (Shen, et al. 2003). A previous study on a set of 25 *Rgh1/Mla* cDNA sequences identified a number of residues/ that have been subject to positive selection, located mainly at the surface-exposed concave side of the deduced LRR solenoid protein structure (Seeholzer, et al. 2010). In contrast, the N-terminal CC domain is mostly invariant among the same set of receptor variants (Seeholzer, et al. 2010). However, this analysis relied predominantly on known *Mla* resistance specificities to *Bgh* in cultivated barley and, therefore, might have underestimated *Rgh1* diversity at the *Mla* locus in wild barley populations.

In this study, we explored *Rgh1* sequence diversity between H. *spontaneum* accessions collected from the Fertile Crescent. In previous studies, due to sequence dissimilarity amongst the three *Rgh* families at *Mla, Rgh1* sequences encoding MLA receptors have been obtained either by PCR with gene-specific primers (Halterman and Wise 2004; Seeholzer, et al. 2010) or DNA gel blot analysis using Rgh1-specific hybridization probes (Wei, et al. 1999; Zhou, et al. 2001). Here, we extracted transcripts belonging to the *Rgh1* family from RNA-Seq data collected from barley leaves. *De novo* transcriptome assembly of 50 wild barley accessions representing nine different wild barley populations (Pankin, et al. 2018) revealed a rich sequence diversity of *Mla* genes that segregate into two major subfamilies. Our findings imply that the divergence of the RGH1/MLA family has been driven by subfamily-specific functionalization to distinct pathogens. Furthermore, interspecies comparison of barley *Mla* and *Mla* orthologs in other cereals identified unique amino acid residues in the wheat Sr33 CC domain, and the importance of these residues was subsequently tested in CC domain modeling and *in planta* functional assays. These natural CC sequence polymorphisms might explain the previously reported differences in tertiary protein structures between the CC domains of barley MLA10 and wheat Sr33 (Maekawa, Cheng, et al. 2011; Casey, et al. 2016).

## Results

### Identification of *Mla* sequences in wild barley accessions

To gain deeper insights into *Mla* diversity in wild barley populations, we used transcriptome sequencing and assembly to identify *Mla* sequences in a set of 50 wild barley accessions that represent nine populations distributed throughout the Fertile Crescent (Pankin, et al. 2018). All accessions were purified by single seed descent to eliminate accession heterogeneity (Pankin, et al. 2018). We included six barley accessions (fig. S1) with already characterized *Mla* resistance alleles to verify that our workflow was able to correctly identify the corresponding gene transcript variants. For each accession, total mRNA was obtained from the first or second leaf at 16-19 hours after inoculation with *Blumeria graminis* f. sp. *hordei* (*Bgh*) conidiospores, as gene expression of *Mla1, Mla6,* and *Mla13* was previously shown to be pathogen-inducible (Caldo, et al. 2004). RNA samples were subjected to paired-end Illumina sequencing, which generated 14-23 Mio read pairs per sample. These RNA-Seq read pairs were then used for *de novo* assembly of transcriptomes for all accessions. Presumptive *Mla* transcripts were extracted from these assemblies by BLAST searches against a database of known *Mla* sequences (for details see Material and Methods).

Using this workflow, we were able to efficiently recover the known *Mla* alleles from all six previously characterized accessions, including those from a complex case in which two *Mla* copies encoding polymorphic MLA variants are present in a single accession (fig. S1). As this initial test verified the suitability of our experimental and bioinformatics pipeline, we next applied this analysis to all wild barley accessions and were able to retrieve *Mla* candidates for all but five of the 50 analyzed accessions (Table S1). Expression levels of the identified *Mla* candidate sequences were variable, ranging from 11 to 300 FPKM (fragments per kilobase of transcript per million mapped reads) for the six previously characterized accessions and from 2 to 140 FPKM for the wild barley accessions (Table S2).

Among the 45 wild barley accessions with putative *Mla* transcripts, in 20 accessions we reliably identified two *Mla* copies and in two accessions we even found three putative *Mla* copies (Table S1). This contrasts with previous findings of a single *Rgh1/Mla* copy in cultivar ‘Morex’ (Wei, et al. 1999; Wei, et al. 2002), suggesting that in wild barley *Rgh1/Mla* has undergone frequent duplication. Five of the 69 identified transcript sequences were excluded from further downstream analysis because in three cases the deduced MLA proteins were C-terminally fused in-frame to a full-length RGH2 family member (i.e. RGH2-MLA) and two other deduced MLA variants lacked the coiled-coil (CC) domain due 185 to truncation of the transcript at the 5’ end. The three RGH2-MLA fusions (FT146-2, FT158 and FT313-2) are 99% identical to each other at the nucleotide level, although their corresponding barley accessions belong to different populations. Moreover, for five accessions a full-length transcript was identified, but due to a premature stop codon the predicted proteins were truncated, lacking part of the NB-ARC domain and the complete LRR. These five sequences were also not included in the phylogenetic analyses unless otherwise stated. Furthermore, we were unable to detect any apparent integrated domains other than the CC, NB-ARC, and LRR among the deduced protein sequences.

### Phylogenetic analysis of MLA sequences

We performed multi-sequence alignment (MSA) and subsequent phylogenetic analysis on the 59 full-length candidate MLA protein sequences retrieved from wild barley together with the 28 previously characterized full-length MLA proteins (Seeholzer, et al. 2010) (excluding truncated MLA38-1) as well as the MLA homologs rmMLA1 from wheat (Jordan, et al. 2011), Sr33 from wheat (Periyannan, et al. 2013), Sr50 introgressed to wheat from *Secale cereale* (Mago, et al. 2015), and the closest homolog to MLA from the more distantly related *Brachypodium distachyon* (XP_014754701.1). The MSA showed that one wild barley accession (FT170) contains the two previously characterized *Mla* alleles *Mla18-1* and *Mla18-2,* whereas two accessions (FT394 and FT355) contain the known *Mla25-1* (fig. 1a, fig. S2). Similarly, in one accession (FT113) one of the two deduced MLA variants differs from already characterized MLA25-1 by only two amino acids. In another accession (FT313) the aligned sequence of one of the two deduced MLA variants is identical to MLA34 but the wild barley sequence shows an extension in the LRR (fig. 1a, fig. S2). These results further confirmed that our method could be used to efficiently recover natural *Mla* variants in wild barley. Our findings indicate that the sequences of at least a few previously published *Mla* alleles have been conserved in extant wild barley populations after their introgression into modern barley cultivars.

**Fig. 1:**
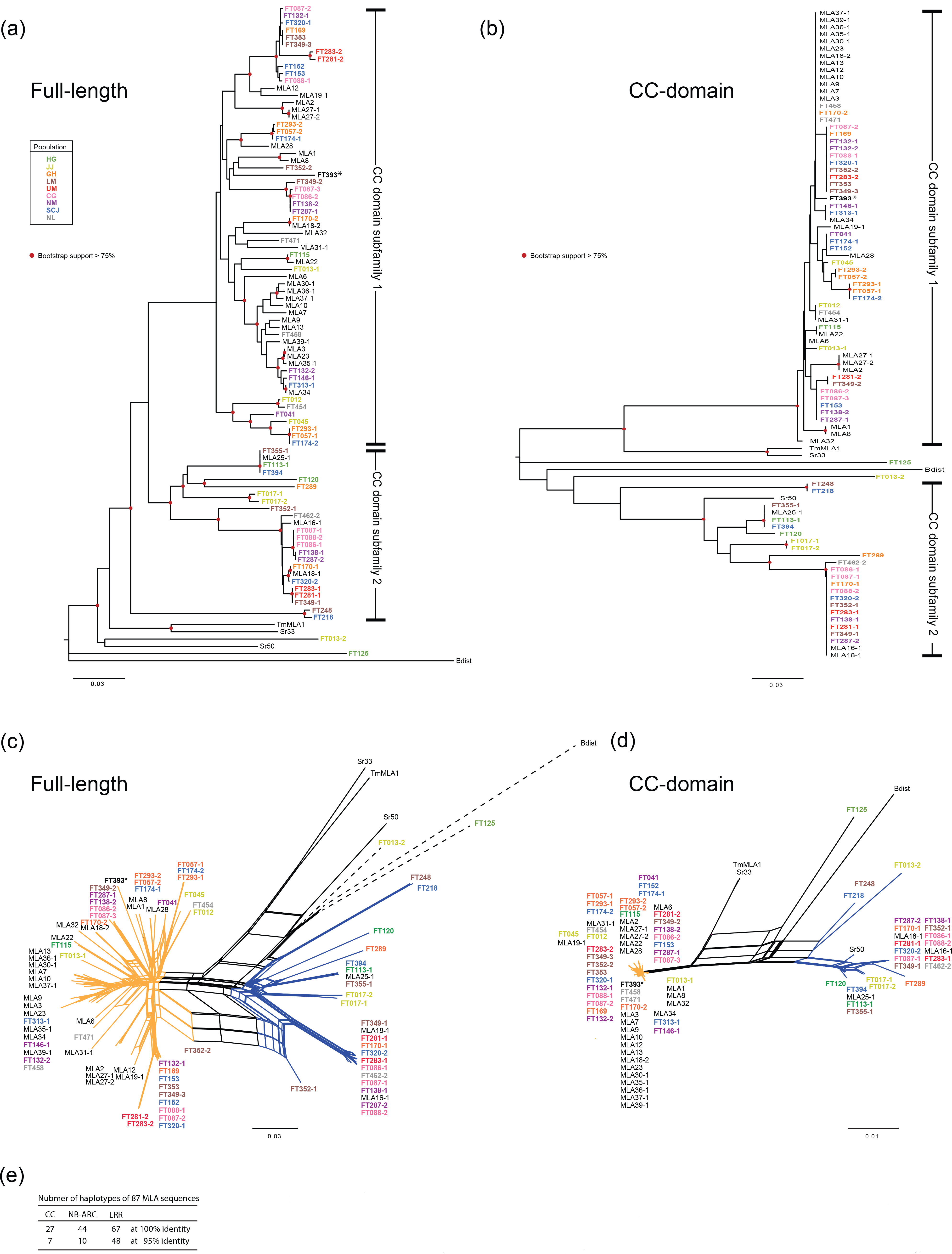
Phylogenetic analysis of 91 MLA and MLA-like protein sequences. (a) Unrooted neighbor-joining (NJ) tree analysis and (c) neighbor-net (NN) analysis of full-length proteins. (b) NJ tree and (d) NN analysis of the CC (coiled-coil) domain (AA 1-151). These analyses were conducted using 28 previously published MLA protein sequences from barley (indicated in black) (Seeholzer et al, 2010), four MLA homologs from other species (indicated in black), and 59 candidate MLA sequences identified in this study from 50 wild barley accessions (colored by population of origin). Only MLA sequences harboring all three domains with more than 895 AA were used in this analysis. (a and b) Red circles mark branches with bootstrap support > 0.75 (500 bootstrap replicates). (c and d) The proteins sequences indicated by orange and blue edges are separately grouped with bootstrap support > 0.95 (1000 bootstrap replicates). The branch length of dashed lines is reduced to half of the actual length. (e) Number of haplotypes of the CC, NB-ARC domains and LRR of 87 barley MLA sequences found in this and previous studies. No *Bgh* resistance activity was detected for MLA16-1, MLA18-1, and MLA25-1 (Seeholzer, et al. 2010). *The population information of FT393 is unavailable.

Overall, the phylogenetic analysis revealed that the newly identified RGH1/MLA proteins from the wild barley accessions do not constitute a distinct clade but are rather distributed across the phylogeny together with known MLA variants conferring *Bgh* resistance (Seeholzer, et al. 2010; Jordan, et al. 2011) (fig. 1a). We conclude that these *Mla* transcripts, retrieved from the wild barley accessions, are derived from naturally occurring *Mla* variants. However, because of markedly lower sequence similarity to the others, two sequences, FT125 and FT013-2, might be derived from an *RGH* family other than *RGH1*. The phylogeny of MLA sequences does not show an obvious relation to the population of origin of the corresponding accessions, while a set of 13 known *Mla* resistance specificities to *Bgh* (e.g. *Mla6, Mla7* and *Mla9*) cluster in the phylogenetic tree (fig. 1a). To crosscheck the results of the neighbor-joining (NJ) phylogenetic analysis, we additionally generated a maximum-likelihood (ML) tree from the same data and compared the two phylogenies (fig. S3). As this comparison showed a good agreement between the two methods, further analyses were performed using the less computationally intensive NJ approach.

When including the five C-terminally truncated sequences in the phylogenetic analysis, we found that four of the five sequences with premature stop codons are identical to each other (fig. S4a). While the four corresponding barley accessions belong to the same UM population, this population also includes accessions harboring different MLA sequences (fig. 1a). Nevertheless, this is the only example in which a wild barley population (i.e. UM population) is seemingly dominated by an invariant RGH1/MLA coding sequence.

A BLAST search of the recently updated barley genome annotations (Mascher, et al. 2017) detected, besides the previously described *Rgh1bcd* copy (Wei, et al. 2002) another potential *Rgh1/Mla* copy (HORVU1Hr1G012190) encoding a protein with a slightly truncated LRR in the genome of cultivar ‘Morex’ (fig. S4b). Although no transcript corresponding to this gene is detected in tissues (http://webblast.ipk-gatersleben.de/barley_ibsc/), an [AT]_n_ microsatellite is found in the third intron, which is commonly found in the six previously isolated *Mla* resistance alleles functioning in *Bgh* resistance (Shen, et al. 2003). Based on the genome of the cultivar ‘Morex’ (Mascher, et al. 2017), the corresponding gene would be located ~ 22 Mb downstream of the previously reported *Mla* locus (Wei, et al. 2002), suggesting that *Rgh1/Mla* family members can be encoded at distant locations in the barley genome.

### Exclusive deployment of one MLA subfamily in modern barley cultivars for *Bgh* resistance

For a more detailed examination of MLA sequence diversity, we focused on the three functional domains of MLA and performed phylogenetic analyses for each domain separately. In agreement with a previous report using mainly barley cultivars (Jordan, et al. 2011), our extended dataset detected two distinct subfamilies in the wild barley accessions, which can be defined by two distinct CC domain haplotypes (fig. 1a, b, fig. S2). Although the bootstrap support for discrimination of the two distinct CC domain subfamilies in the neighbor-joining tree is not very high, neighbor-net analysis provided supporting evidence with very high bootstrap support (> 95%: fig. 1 c and d). This result suggests that the underlying evolutionary history of *Rgh1/Mla* is not tree-like. Furthermore, we found a striking pattern of CC sequence diversification, characterized by differential occupancy of charged or noncharged amino acids in subfamily 1 and 2 (indicated by arrows; fig. S5). These charge alterations ultimately distinguish the two MLA/RGH1 subfamilies. Thirty-nine (61%) of the 64 MLA sequences from wild barley (including the five C-terminally truncated proteins) belong to CC domain subfamily 1 and 39% to subfamily 2 (Table S1). Within the 64 novel sequences and 29 known MLA sequences, we can distinguish 27 sequence haplotypes for the CC domain, of which 16 belong to subfamily 1 and nine to subfamily 2 (fig. 1e, Table S3). Notably, all known MLA variants conferring resistance to *Bgh* belong exclusively to subfamily 1. Furthermore, the most common CC haplotype (haplotype CC01) for the MLA variants providing *Bgh* resistance (13 of 25 variants; Table S3) is found only in three wild barley accessions (FT170-2, FT458 and FT471). These data suggest that the CC haplotype in subfamily 1 might confer resistance to *Bgh* in agricultural settings and that this haplotype has predominantly been deployed in modern barley cultivars.

As for the full-length and CC domain sequences, NJ phylogenetic and neighbor-net analyses based on the NB-ARC domain and LRR showed that many clades contain the reported MLA sequences and sequences from wild accessions (fig. 2a-d). We distinguished a total of 44 sequence haplotypes for the NB-ARC domain (fig. 1e). Among them, NB-ARC domains of 12 MLA variants conferring *Bgh* resistance constitute the most common haplotype together with those of FT458 and FT313-1 (fig. 2a, c). Whilst the clear separation into the two MLA subfamilies is partly retained for the NB-ARC domain (fig. 2c), the LRR does not contribute to the subfamily grouping (fig. 2d). Moreover, pair-wise comparisons of the phylogenies for the three domains show that while certain smaller phylogenetic groups might be conserved across domains, major discrepancies can be detected especially between the LRR phylogeny and the NB-ARC and CC phylogenies (fig. S5). This suggests that the three domains, especially the LRR, have evolved largely independently from each other.

**Fig. 2:**
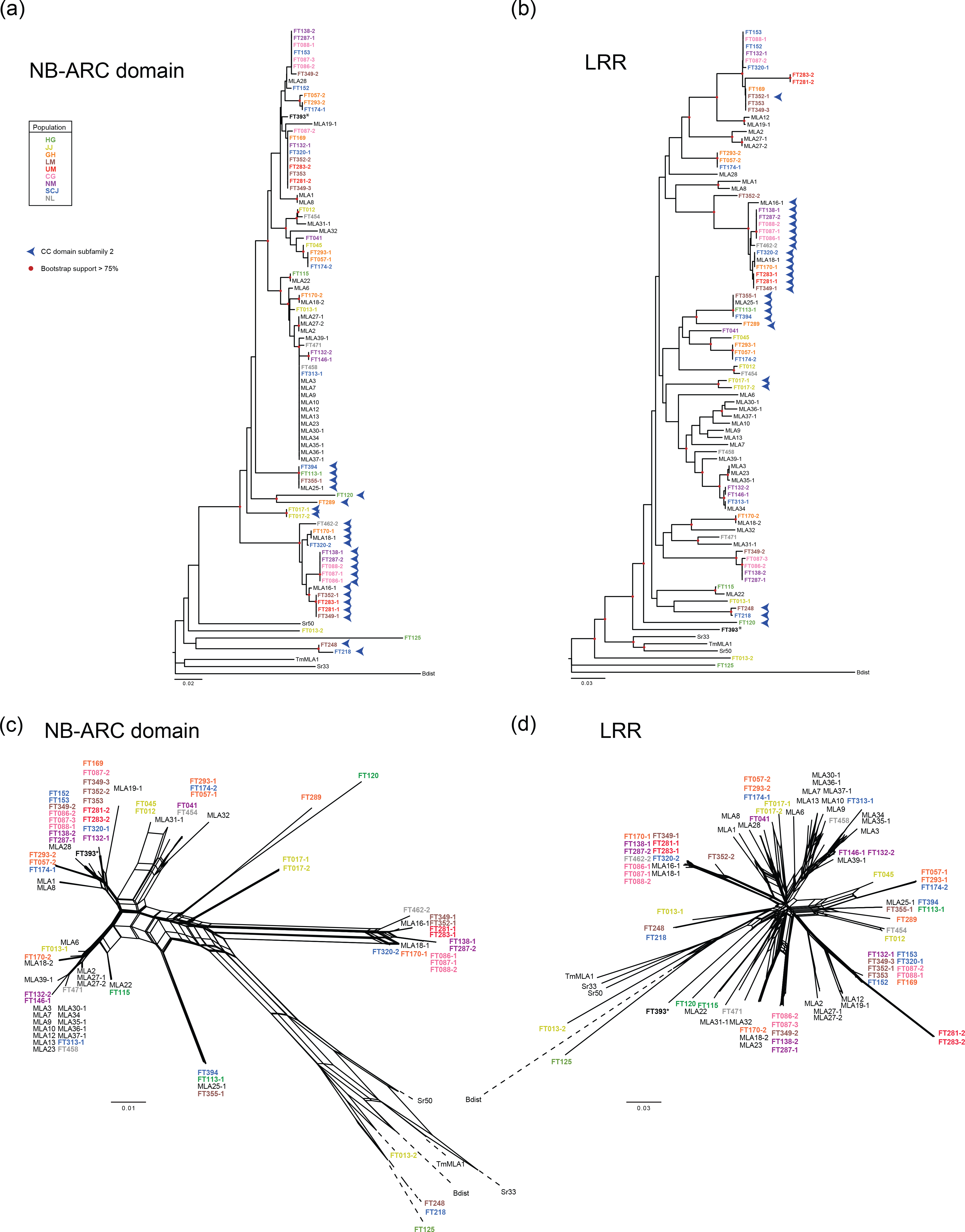
Phylogenetic analysis of the MLA protein domains. (a) NJ tree analysis and (c) NN analysis of the MLA NB-ARC (nucleotide-binding adaptor shared by APAF-1, R proteins, and CED-4) domain (AA 180-481). (b) NJ tree analysis and (D) NN analysis of the MLA LRR (leucine-rich repeat region) (AA 1490-end). These analyses were conducted using 28 previously published MLA protein sequences from barley (indicated in black) (Seeholzer, et al. 2010), four sequences of MLA homologs in other species (indicated in black), and 59 candidate MLA sequences that were identified in this study from 50 wild barley accessions (colored by population of origin). Only MLA sequences harboring all three domains with more than 895 AA were used in this analysis. Red circles mark branches with bootstrap support > 0.75 (500 bootstrap replicates). Blue arrowheads indicate members of CC domain subfamily 2 shown in fig. 1. The branch length of dashed lines (c) and (d) is reduced to 10% and half of the actual length, respectively. *The population information of FT393 is unavailable.

### The majority of sites under positive selection locate to the LRR

To estimate MLA sequence diversity, we excluded the two most divergent candidate sequences (FT125 and FT013-2), as we could not exclude that these represent RGH proteins other than RGH1. Based on the remaining 57 full-length candidate MLA sequences and the 28 previously published full-length MLA variants we observed an overall nucleotide diversity (π) of 0.068. Sequence diversity within the wild accessions (π = 0.072) is slightly higher than in the previously published MLA variants (π = 0.055) and sequence diversity is also higher in subfamily 2 (π = 0.060) compared to subfamily 1 (π = 0.045; Table S4).

The pair-wise comparisons of the phylogenies for the three MLA domains (fig. S5) demonstrate highly domain-specific sequence selection within a single gene. We thus performed dedicated statistical analyses on the coding sequences of the 85 full-length MLA sequences to identify sites under episodic (MEME; Mixed Effects Model of Evolution) or pervasive (FUBAR; Fast Unconstrained Bayesian AppRoximation) positive selection. The former analysis was included to also allow detection of positive selection sites acting on a subset of branches in a phylogeny. In accordance with previous observations (Seeholzer, et al. 2010), the majority of sites under positive selection, whether pervasive or episodic, are located in the LRR (fig. 3a). Using this complete dataset, we also observed a number of sites under episodic positive selection (*p* < 0.1) in the CC domain, which mostly involve residues that differ between the two subfamilies (fig. 3a). Accordingly, when we performed separate analyses for the two subfamilies, we observed that within each subfamily only very few sites under positive selection are in the CC or NB-ARC domains (fig. 3b,c). In these separate analyses for both subfamilies, sites under pervasive positive selection (posterior probability > 0.95) are detected almost exclusively in the LRR, and many of these sites in the LRR seem to be under positive selection within both subfamilies as well as in the complete set (fig. 3). The presence of additional sites under episodic positive selection in subfamilies 1 and 2 suggests distinct selection pressures acting on a few branches of both subfamilies (fig. 3b, c). The location and clustering of sites under positive selection for subfamily 1 is largely conserved between the sequences from wild barley and known MLA resistance specificities to *Bgh* (fig. S7a, b). Among 18 sites for the wild accessions and 16 sites for known MLA resistance specificities to *Bgh* that are located in the β-strand motifs in the LRR, 13 sites are shared (fig. S7c-f).

**Fig. 3:**
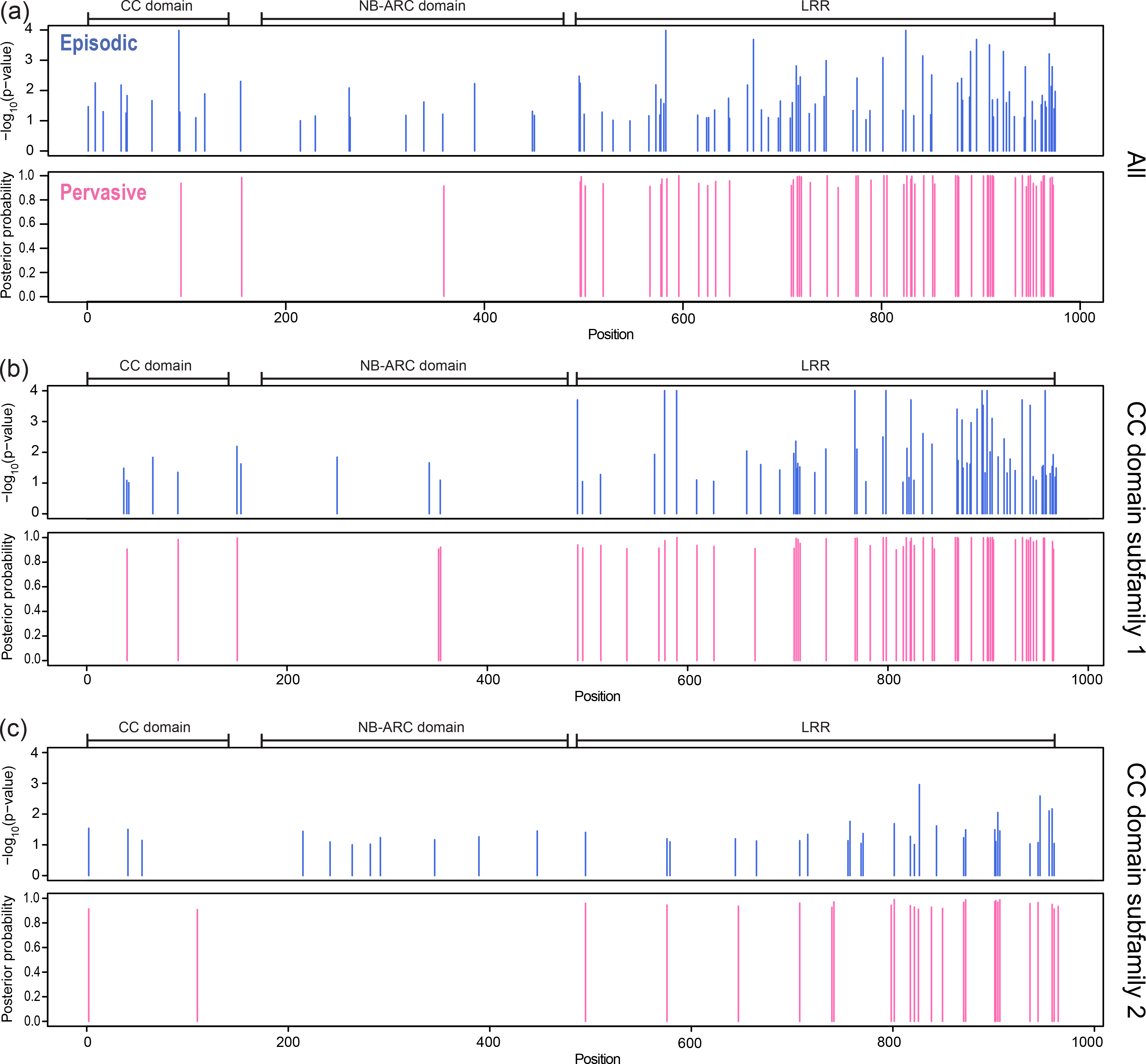
Identification of positively selected sites in previously known and newly identified candidate MLA cDNAs. (a) Sites under episodic (upper panel; blue bars) or pervasive (lower panel: pink bars) positive selection in a set of 85 known and newly identified MLA cDNAs. (b) Sites under episodic (upper panel; blue bars) or pervasive (lower panel: pink bars) positive selection in 61 known and newly identified MLAs carrying a CC domain belonging to subfamily 1. (c) Sites under episodic (upper panel; blue bars) or pervasive (lower panel: pink bars) positive selection in 24 known and newly identified MLAs carrying a CC domain belonging to subfamily 2. Only full-length MLA sequences (at least 895 AA) from barley were included in these analyses. To test for episodic selection, we used MEME and judged all sites with a p value below 0.1 to be under positive selection. To test for pervasive selection, we additionally used FUBAR and judged all sites with a posterior probability above 0.95 to be under positive selection. AA 1-151: CC domain; AA 180-481: NB-ARC domain; AA 490-end: LRR.

### The subfamily 2 CC domain is functional in cell death

The ability of RGH1 members from subfamily 2 to confer resistance to *Bgh* remains unclear, as no *Bgh* resistance activities were detected for MLA16-1, MLA18-1, and MLA25-1 (Seeholzer, et al. 2010; Jordan, et al. 2011). This raises the question of whether subfamily 2 MLA receptors can generally act as disease resistance proteins. Our phylogenetic analyses of MLA CC domains allowed us to detect a close relationship between Sr50 (conferring resistance to *Puccinia graminis* f.sp. *tritici*; *Pgt* in wheat) and subfamily 2 MLAs (fig. 1b, d). Overexpression of MLA10 representing the haplotype CC01 (Table S3), Sr33 and Sr50 N-terminal CC domains initiates downstream signaling events that are comparable to the signaling mediated by the corresponding full-length receptors, including the execution of cell death (Maekawa, Cheng, et al. 2011; Casey, et al. 2016; Cesari, et al. 2016; Jacob, et al. 2018). We thus examined whether subfamily 2 MLA CC domain variants are capable of triggering cell death *in planta*. Overexpression of the CC domain of FT394, one of the closest barley variants of wheat Sr50, was able to induce cell death in *Nicotiana benthamiana* leaves (fig. 4), suggesting that *Mla* alleles in subfamily 2 encode receptors with cell death activity. Although the FT394 CC variant is 93% and 67% identical to the CC domains of Sr50 and MLA10, respectively, the time of onset and confluence of the necrotic lesions induced by FT394 CC and MLA10 CC were comparable, whereas lesions elicited by the expression of Sr33 and Sr50 CC domains remained invariably patchy.

**Fig. 4:**
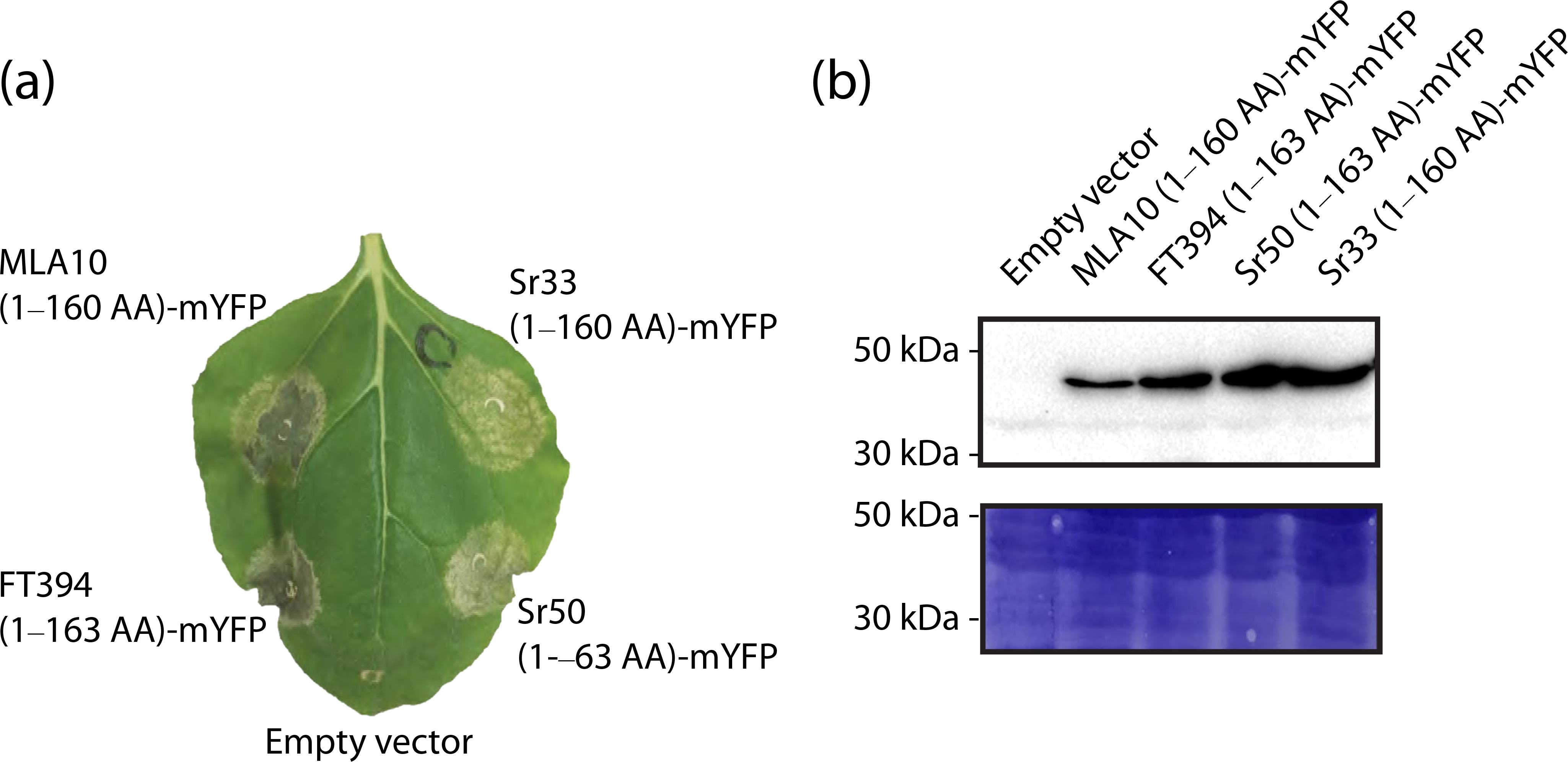
CC domains of subfamily 2 MLA FT394 exhibits cell death activity. (a) *Nicotiana benthamiana* plants were transiently transformed to express the CC domains of MLA10_1-160 amino acids (AA), FT394_1-163AA, Sr50_1-163AA, and Sr33_1-160AA, each fused C-terminally to monomeric YFP, or empty vector control; the picture was taken three days post infiltration. (b) Immunoblot analysis corresponding to (a). Transformed leaf tissue was harvested 24 hours post infiltration. Proteins were analyzed after gel electrophoresis and western blotting with an anti-GFP antibody.

### The Sr33 CC domain harbors unique amino acid substitutions

In contrast to Sr50, Sr33 is assigned to a clade of the CC domain together with *TmMLA1* (fig. 1b, d). Notably, interspecies comparison of MLA or MLA-like sequences identified unique amino acid polymorphisms in the Sr33 and TmMLA1 CC-domains (fig. 5a). The 21^st^ positions of the CC domains of MLA functioning in *Bgh* resistance, MLA from wild barley and homologs of some Triticeae family members are generally occupied by aspartate (D) or glutamate (E), but glycine (G) occupies the corresponding positions in Sr33 and TmMLA1. The same substitution is found in an accession of rye (*Secale cereale*) (fig. 5a). The structure of the Sr33 CC domain (6-120 amino acids) adopts a monomeric four-helix bundle conformation, while the structure of the MLA10 CC domain (5-120 amino acids) is arranged in an antiparallel homodimer that adopts a helix-loop-helix fold (Maekawa, Cheng, et al. 2011; Casey, et al. 2016; Fig, 5B). Intriguingly, in the corresponding structures the 21^st^ position of MLA10 locates to the middle of the alpha helix, while in Sr33 this position with an adjacent valine corresponds to a loop region of the CC domain (fig. 5b, fig. S8a). These amino acid differences might account for the differences in tertiary protein structure between the CC domains of MLA10 and Sr33.

**Fig. 5.**
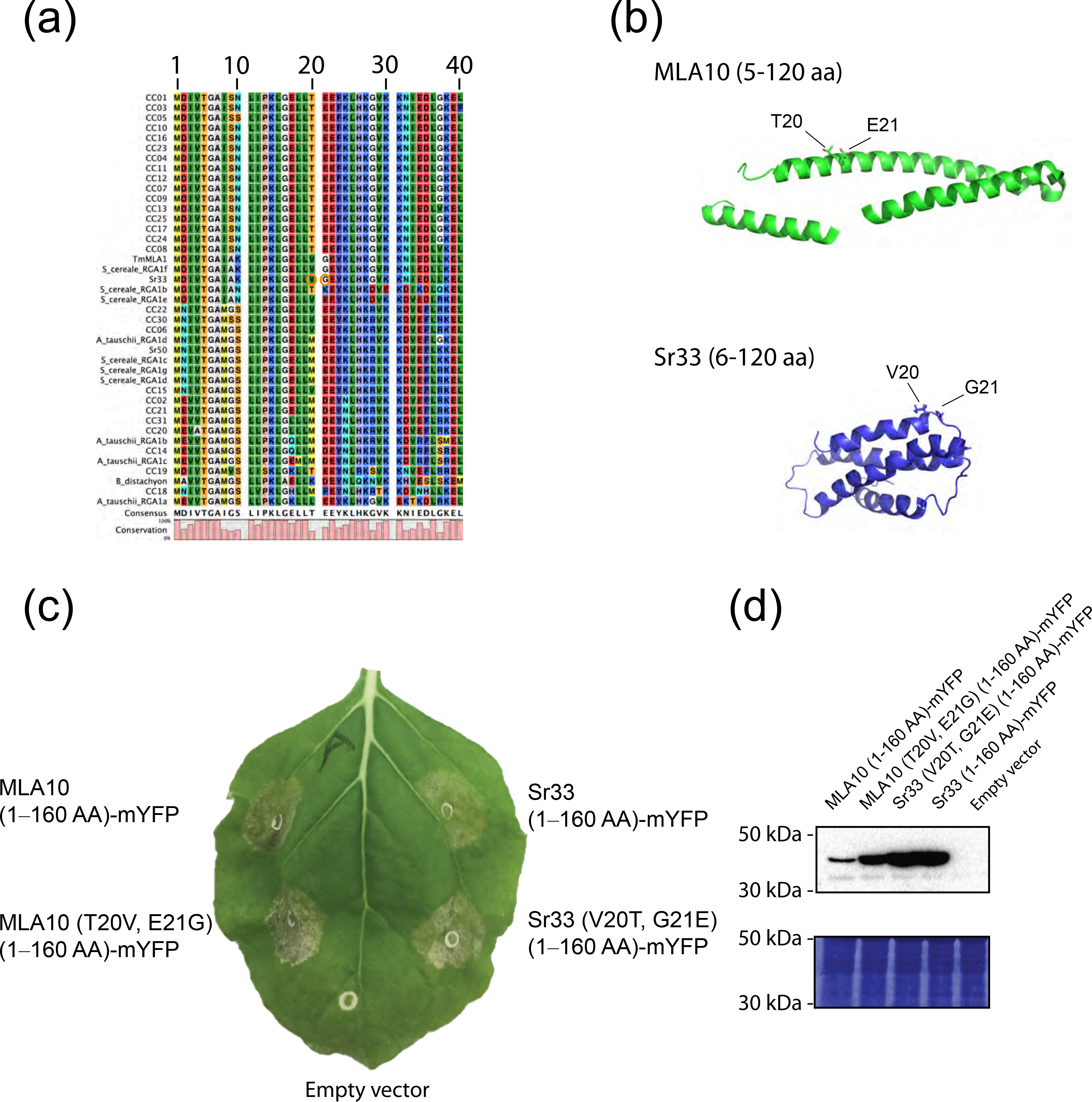
Sr33 carries unique amino acid substitutions in a loop region of the coiled-coil (CC) domain. Amino acid sequence alignment of the CC variants (1-40 amino acids (AA), see Table S3b) of MLA and MLA orthologs. The unique residues in the Sr33 CC domain are indicated by orange circles. (b) The solved tertiary protein structures of MLA10 (PDB ID 3QFL) and Sr33 (PDB ID 2NCG) CC domains. A protomer of the CC domain dimer of MLA10 is shown. (c) The unique amino acids do not contribute to the intensity of cell death in *Nicotiana benthamiana. N. benthamiana* plants were transiently transformed to express the CC domains of MLA10_1-160AA, MLA10_1-160AA (T20V, E21G), Sr33_1-160AA, and Sr33_1-160AA (V20T, G20E) each fused C-terminally to monomeric YFP, or EV and a picture was taken three days post infiltration. (d) Immunoblot analysis corresponding to (c). Transformed leaf tissue was harvested 24 hours post infiltration. Proteins were analyzed after gel electrophoresis and western blotting with an anti-GFP antibody.

To examine the role of these two amino acids in structural folding, we simulated secondary structures of wild-type and mutated CC domains of MLA10 and Sr33 using PSIPRED (Buchan, et al. 2013). For simplicity, we used the first 40 amino acids for the two following analyses. MLA10 (wild-type) and the mutated MLA10 (T20V E21G) that carries the Sr33-type residues were predicted to be structurally similar, while Sr33 (wild-type) was predicted to contain an additional loop compared to the MLA10 variants. Notably, this loop was no longer predicted to form in the Sr33 (V20T G21E) mutant carrying the MLA10-type residues (fig. S8b). We additionally assessed the impact of each single mutation using STRUM (Quan, et al. 2016). Both T20V and E21G substitutions in MLA10 were predicted to decrease the protein stability, while V20T but not G21E was predicted to stabilize the Sr33 structure. These data suggest that the two unique residues (or either one of the two) in Sr33 could destabilize the helix fold in its CC structure and, as a consequence, the overall CC structure of Sr33 might differ from the CC domain of MLA10.

We then experimentally tested cell death mediated by the CC domains of MLA10 (wild-type) and a site-directed mutant form, MLA10 (T20V E21G), carrying the Sr33-type residues. Necrotic lesions in *N. benthamiana* leaves elicited by these two CC domain variants were comparable in size and macroscopic appearance. In reciprocal experiments, lesions elicited by the CC domains of Sr33 (wild-type) and mutant Sr33 (V20T G21E) carrying the MLA10-type residues were also comparable (fig. 5c). Thus, these amino acid sequence polymorphisms in the two CC domains do not abrogate cell death mediated by both NLRs.

## Discussion

### Divergence of *Mla* subfamilies in wild barley populations

We revealed a rich sequence diversity of the *Rgh1/Mla* gene family in wild barley accessions representing nine populations distributed throughout the Fertile Crescent of the Middle East. In addition, in wild barley *Rgh1/Mla* has undergone frequent gene duplication (Table S1), necessitating a revision of the current view that *Mla* recognition specificities to *Bgh* represent alleles of a single gene (Shen, et al. 2003). We found no clear evidence for geographic isolation and population structure of particular *Mla* sequence clades. Similarly, a genome-wide survey of *NLR* genes in *A. thaliana* revealed no evidence for regional selection of particular disease resistance genes (Bakker, et al. 2006). The observed similar ratio of *Mla* subfamily 1 and 2 members among individual wild barley accessions (37 and 27, respectively; Table S1) was unexpected and suggests that a balancing selection mechanism maintains the two subfamilies in the host populations. Pathogen selection pressure is likely driving the observed sequence diversification at least within subfamily 1 because all known *Mla* resistance specificities to *Bgh* belong to this subfamily (Seeholzer, et al. 2010; Jordan, et al. 2011). Compared to the CC domain, sequences of the LRR do not group into two subfamilies (fig. 2d). This contrasting pattern indicates that selection regimes differ between the CC and LRR domains that might involve recombination events that swap LRRs between two *Rgh1/Mla* alleles in the two different subfamilies. A QTL analysis on disease resistance in the field, conducted in a cross derived from the barley cultivar ‘Carola’ and the wild barley accession FT394 that carries only one subfamily 2 member (Table S1), did not reveal any powdery mildew resistance mediated by the *Mla* locus (Wang 2005). In addition, the *Mla* locus did not associate with any of 13 tested agronomic traits in a population derived from a cross between barley cultivar ‘Apex’ and FT394, suggesting that this *Mla* allele does not affect agronomic performance in the field (Pillen, et al. 2003). Thus, the exclusive deployment of subfamily 1 members in modern barley cultivars for *Bgh* resistance is probably not due to any negative impacts on agronomical performance associated with introgression of subfamily 2. Conservation of NLR motifs needed for immune receptor function (fig. S2) (Tameling, et al. 2006; Rairdan, et al. 2008), together with positively selected sites in the LRR among subfamily 2 members (fig. 3c), implies a similar functional diversification of subfamily 2 NLRs in response to barley pathogen(s). Whether subfamily 2 NLRs confer disease resistance to avirulence genes present in yet uncharacterized *Bgh* populations or other pathogens remains to be tested. The presence of both subfamily 1 and 2 members in genomes of several accessions (14 out of 50 accessions; Table S1) also predicts non-redundant immune functions of two subfamilies.

The CC domain sequences of barley subfamily 2 members are closely related to that of Sr50, which is encoded by an *Mla* ortholog originating from rye (*Secale cereale*) and was introgressed into wheat for resistance against the stem rust fungus *Puccinia graminis* f. sp. *tritici* (*Pgt;* (Mago, et al. 2015). Rye and wheat diverged from each other approximately 3−4 Mya and barley diverged from wheat 8−9 Mya (Krattinger and Keller 2016). Closer inspection of the phylogenetic tree of MLA CC domains indicates that subfamily 2 CC is evolutionarily maintained, at least in rye and barley (fig. 1b and d). In contrast, the presence of subfamily 1 members appears to be restricted to barley (fig. 1b and d). The CC domains of both barley MLA subfamilies are capable of cell death initiation, further supporting our hypothesis that members belonging to subfamily 2 also encode NLRs mediating innate immunity. The MLA subfamilies are mainly defined by two major haplotypes of the N-terminal CC signaling domain rather than the C-terminal LRR that has been shown to determine AVR recognition specificities in *Bgh* resistance (fig. 1 and 2; Shen, et al. 2003). What evolutionary mechanism has maintained the two barley MLA subfamilies in the host population? To date, only MLA subfamily 1 members have been shown to confer immunity to the barley powdery mildew fungus (Seeholzer, et al. 2010; Jordan, et al. 2011). One possible explanation is that *Bgh* has overcome immune signaling mediated by the CC of subfamily 2 members. For example, the effectors of the *Pgt* stem rust pathogen can suppress cell death mediated by the Sr50 CC domain when overexpressed (Chen, et al. 2017). However, overexpression of a barley MLA chimera containing CC and NB-ARC domains of MLA25-1, a subfamily 2 member of unknown function, fused to the LRR of barley MLA1, a subfamily 1 member, retains MLA1-dependent resistance to *Bgh* (Jordan, et al. 2011). This finding might suggest that subfamily 1 and 2 CC domains can be interchanged without affecting effector recognition. Together, this demonstrates that subfamily 2 CC and NB-ARC domains can be functional in *Bgh* resistance when overexpressed but does not exclude the possibility that in wild barley populations, with native *Mla* gene expression levels, immunity to *Bgh* mediated by the CC of subfamily 2 members is inefficient in limiting pathogen reproduction.

Consistent with extensive CC diversification in the phylogeny of MLA subfamilies 1 and 2, the corresponding sequence alignment reveals 22 residues that are differentially occupied by charged or non-charged amino acids in the two subfamilies (indicated by arrows fig. S5). These charged residues are likely surface-localized (Maekawa, Cheng, et al. 2011; Casey, et al. 2016), thereby altering the surface charge distribution of the signaling module. Interestingly, the CC domains of MLA10 (subfamily 1), but not MLA18-1 and MLA25-1 (both subfamily 2), interact with barley WRKY transcription factors that repress immune responses to powdery mildew fungi (Shen, et al. 2007; Jordan, et al. 2011). The N-terminal 46 amino acids of the barley MLA10 CC domain, including three of the aforementioned 13 charge alterations, are critical for this interaction (Shen, et al. 2007). Therefore, whilst the CC of MLA subfamily 1 is predicted to de-repress immune responses via direct interaction with these WRKYs (Shen, et al. 2007), the CC of MLA subfamily 2 members might interact with other host proteins and interfere with the same transcriptional machinery or another cellular process conferring immunity and cell death.

### Does CC signaling domain diversification suggest diversification of CC signaling pathways?

Barley MLA10 was shown to localize to both the nucleus and the cytoplasm and experiments involving enforced mis-localization of the receptor suggest cell compartment-specific bifurcation of receptor-mediated cell death and disease resistance signaling in the cytoplasm and nucleus, respectively (Shen, et al. 2007; Bai, et al. 2012). Different CC structural folds have been reported for sequence-related barley MLA10 and wheat Sr33 (Maekawa, Cheng, et al. 2011; Casey, et al. 2016). This has given rise to a model in which the two structures represent closed “off-state” and open “on-state” conformations of the corresponding full-length NLRs, possibly linked to receptor oligomerization (El Kasmi and Nishimura 2016). Proteins or protein modules that can adopt more than one native folded conformation are more common than previously thought and are designated metamorphic or fold-switching proteins (Murzin 2008; Porter and Looger 2018). Our interspecies comparison identified unique amino acid polymorphisms between barley MLA and wheat Sr33 CC domains (T20E21 and V_20_G_21_, respectively; fig. 5A). Directed substitutions of T_20_E_21_ to V_20_G_2_i in the barley MLA10 CC do not significantly affect the capacity of the N-terminal domain alone to trigger cell death in *N. benthamiana* and, similarly, the reciprocal substitutions in the wheat Sr33 CC did not impair its capacity for cell death activation. If the unique natural CC amino acid polymorphisms directly contribute to the reported structural differences of MLA10 and Sr33 CC modules, one would have expected that one of the mutant CC variants would show weakened cell death in the *N. benthamiana* leaf assay. The extent of cell death was macroscopically indistinguishable between wild-type and CC mutant variants, showing that both structures are cell death signaling-competent forms. We assume that in *N. benthamiana* leaves the confluent cell death phenotype linked to MLA10 CC expression compared to the characteristic cell death patches elicited by Sr33 CC expression reflects immune responses of differing strengths triggered by the respective CC modules. Given that both characteristic cell death phenotypes were retained upon expression of the tested CC mutants (fig. 5), other amino acid polymorphisms between MLA10 and Sr33 CC, including the predicted differential surface charge distribution and differential interactions with other host signaling components, must account for the observed macroscopic differences in cell death. In this model, the equilibrium between receptor ‘on’ and ‘off’ states might influence the ratio of nuclear to cytoplasmic MLA-mediated defense outputs through differential interactions with host signaling components. This could explain why enforced cytoplasmic receptor localization is sufficient to induce stem rust resistance in transgenic Sr33-expressing wheat (Cesari, et al. 2016), whereas the nuclear receptor pool of barley MLA10 is necessary to induce *Bgh* resistance (Shen, et al. 2007). Similarly, divergence of the barley RGH1/MLA signaling domain in wild barley might indicate haplotype-dependent immune responses that are maintained to control distinct pathogen populations.

### The LRR as pathogen recognition determinant

The location and clustering of sites under positive selection for subfamily 1 is generally conserved between the MLA sequences from wild barley and functionally validated receptor variants conferring resistance to *Bgh* (fig. 3, fig. S7). Notably, the majority of positive selection sites are found on the surface of the predicted concave side of the LRR solenoid and these sites are mostly shared between the natural MLA variants (fig. S7). This implies that the previously reported sites under positive selection, which were detected among a set of MLA variants conferring *Bgh* resistance (Seeholzer, et al. 2010), closely reflect the evolutionarily pressure acting on this receptor region in wild barley populations. The positively selected sites are mainly located between leucine-rich repeats 10 to 15 in MLA subfamilies 1 and 2. Wheat Pm3 is predicted to contain 28 leucine-rich repeats with positively selected sites located mainly between repeats 19 and 28 (Krattinger and Keller 2016). Recent data from our lab suggest that MLA receptors directly recognize sequence-unrelated *Bgh* avirulence effectors (Saur and Bauer et al., unpublished data). Thus, the clustering of positively selected amino acid residues in the C-terminal leucine-rich repeats of both MLA receptor subfamilies might define contact residues for direct effector interactions that do not compromise conformational change(s) of the full-length receptor from the off-state to the on-state. The relevance of positive selection sites for direct interactions with avirulence proteins has been shown for a subset of L resistance alleles in flax conferring immunity to the rust pathogen *Melampsora linii* (Dodds, et al. 2006; Wang, et al. 2007).

### Mining wild barley germplasm for novel *Mla* resistance specificities

In wheat, the *Pm3* locus is the main source of genetically encoded disease resistance against the wheat powdery mildew fungus, B. *graminis* f. sp. *tritici (Bgt;* Krattinger and Keller 2016). Similar to barley *Mla,* 17 allelic variants of the *Pm3* gene have been identified in wheat populations that determine isolate-specific immunity to *Bgt* carrying matching AVR genes (Krattinger and Keller 2016). Although *Pm3* is widely deployed in domesticated hexaploid wheat (*Triticum aestivum),* in *T. dicoccoides,* the progenitor of most cultivated wheat species, high presence-absence variation of polymorphism in the *Pm3* gene was observed as well as a low sequence diversity (61% of 208 accessions lacked the *Pm3* gene; (Srichumpa, et al. 2005; Sela, et al. 2014)). This has been explained by a potential maintenance cost of *Pm3* in *T. dicoccoides* in the absence of pathogen selection pressure. Consequently, it is thought that the observed functional diversification in *Pm3* has occurred primarily after wheat domestication ~ 10,000 years ago (Srichumpa, et al. 2005; Sela, et al. 2014). This contrasts with the detection of *Mla* transcripts encoding sequence-diversified full-length MLA immune receptors in at least 40 out of 50 tested wild barley accessions. This and the fact that at least some characterized *Mla* recognition specificities to *Bgh* in cultivated barley are derived from wild barley or barley landraces might help to explain why 25 functionally validated *Mla* recognition specificities to *Bgh* exhibit on average > 91% sequence identity, whereas 17 deduced allelic wheat Pm3 receptors share > 97% sequence identity (Seeholzer, et al. 2010). Accordingly, continuous *Bgh* selection pressure and a much longer evolutionary time span has been available for *Mla* functional diversification in wild barley, and this could account for the greater sequence diversity of MLA compared to Pm3 NLRs. Several novel natural *Mla* sequences from wild barley are distributed across the *Mla* phylogenetic tree, with functionally validated *Mla* resistance specificities to *Bgh* confined to subfamily 1. A conspicuous clustering of *Mla* resistance specificities is seen in a sublineage of subfamily 1 containing 13 known resistance specificities, including *Mla6, Mla7, Mla9, Mla10,* etc. (fig. 1). The identification of four novel *Mla* sequence variants from wild barley belonging to this sublineage makes these receptor variants prime candidates for future targeted disease resistance assays. Such experiments could be used to examine whether the receptor variants exhibit overlapping recognition specificities with known MLA receptors or to detect novel *Bgh AVR_A_* genes.

## Supplementary figures

**fig. S1:**
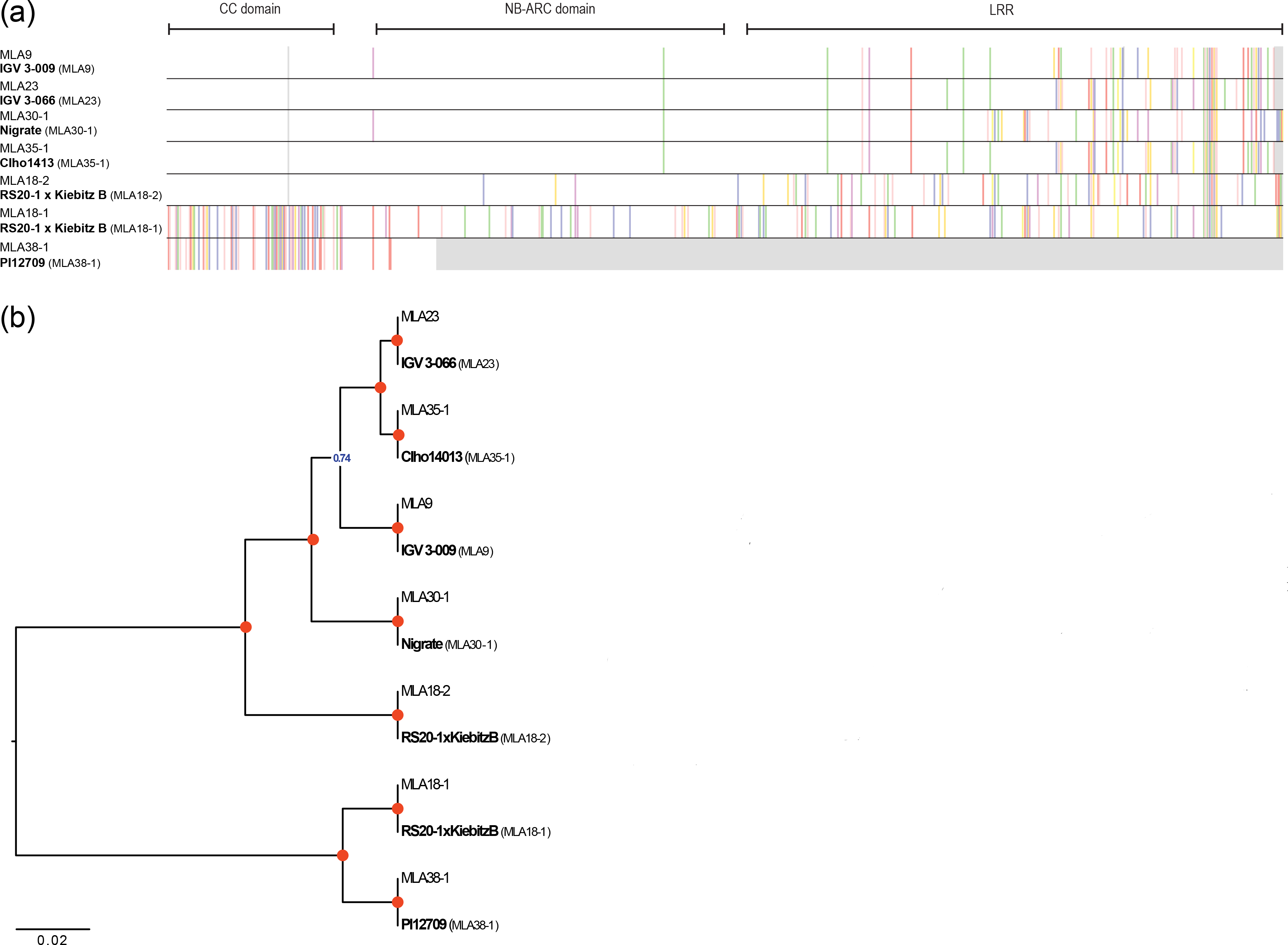
Validation of MLA identification workflow through recovery of known MLA sequences. (a) Amino acid (AA) sequence alignment of seven known MLA alleles and the corresponding sequences as obtained by our recovery workflow starting from RNA-sequencing data of the indicated barley lines. For visualization, an AA consensus sequence was obtained from 28 previously published full-length barley MLA sequences (Seeholzer, et al. 2010) and for each MLA polymorphic residues relative to this consensus are highlighted in color (gray color indicates alignment gaps). The three domains of MLA are indicated at the top. (b) Unweighted Pair Group Method with Arithmetic mean tree analysis of the 14 MLA sequences. Tree calculation was performed using the pairwise deletion option. Red circles mark branches with a bootstrap support of 1 (1,000 bootstrap replicates).

**fig. S2:**
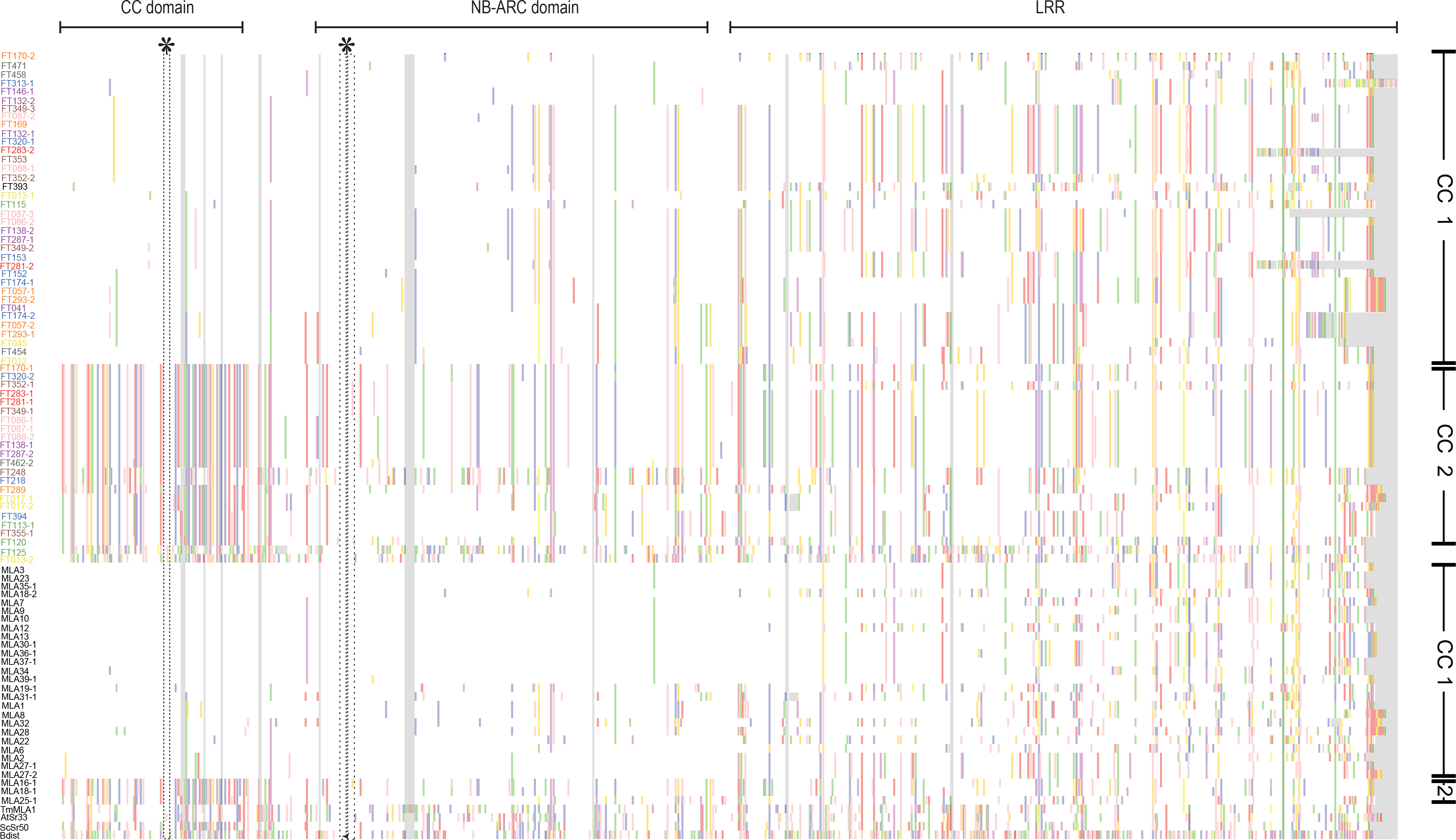
Amino acid (AA) sequence alignment of 91 full-length MLA protein sequences. The AA alignment includes full-length sequences of 28 previously published MLA protein sequences from barley (indicated in black) (Seeholzer et al, 2010), four MLA homologs from other species (indicated in black), and 59 candidate MLA sequences identified in this study from 50 wild barley accessions (colored by population of origin). For visualization, an AA consensus sequence was obtained from the 28 previously published barley MLA sequences and polymorphic residues in each MLA relative to this consensus are highlighted in color (gray color indicates alignment gaps). The three domains of MLA are indicated at the top. Asterisks mark the EDVID motif (Rairdan, et al. 2008) in the coiled-coil (CC) domain and the Walker A motif (Tameling, et al. 2006) with the p-loop in the nucleotide-binding (NB) domain. Only full-length MLA sequences (at least 895 AA) were included in this analysis.

**fig. S3:**
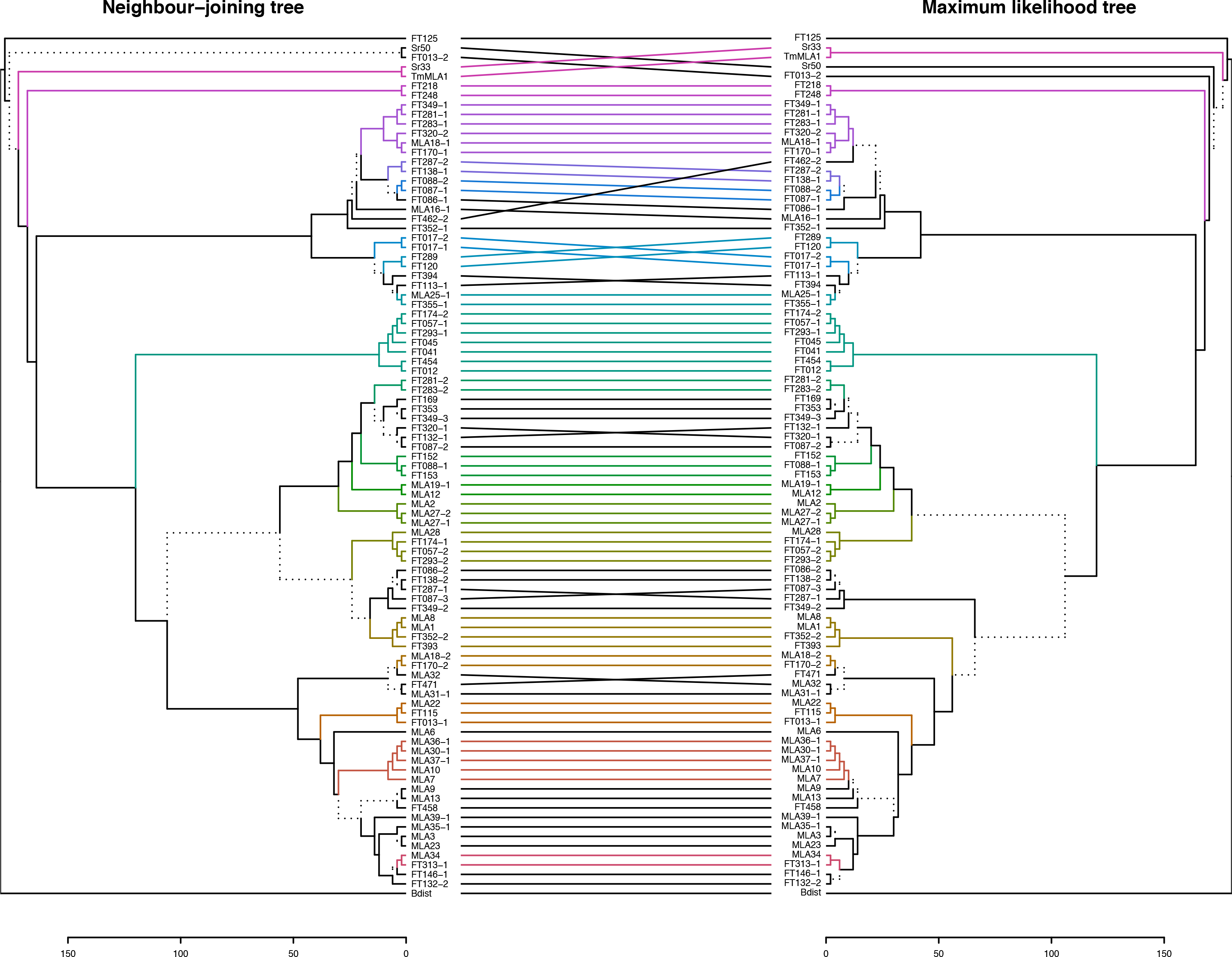
Comparison of neighbor-joining (NJ) and maximum-likelihood (ML) trees obtained from 91 full-length MLA protein sequences. Trees include full-length amino acid (AA) sequences of 28 previously published MLAs from barley (Seeholzer, et al. 2010), four MLA homologs from other species, and 59 candidate MLA sequences identified in this study from 50 wild barley accessions. All positions containing gaps or missing data were omitted from the tree calculations (complete deletion option). A tanglegram was created from the two trees using the R package ‘dendextend’, in which lines connect the same isolates and colors represent sub-trees conserved between the two methods. Branches shown as dashed lines indicate differences between the trees.

**fig. S4:**
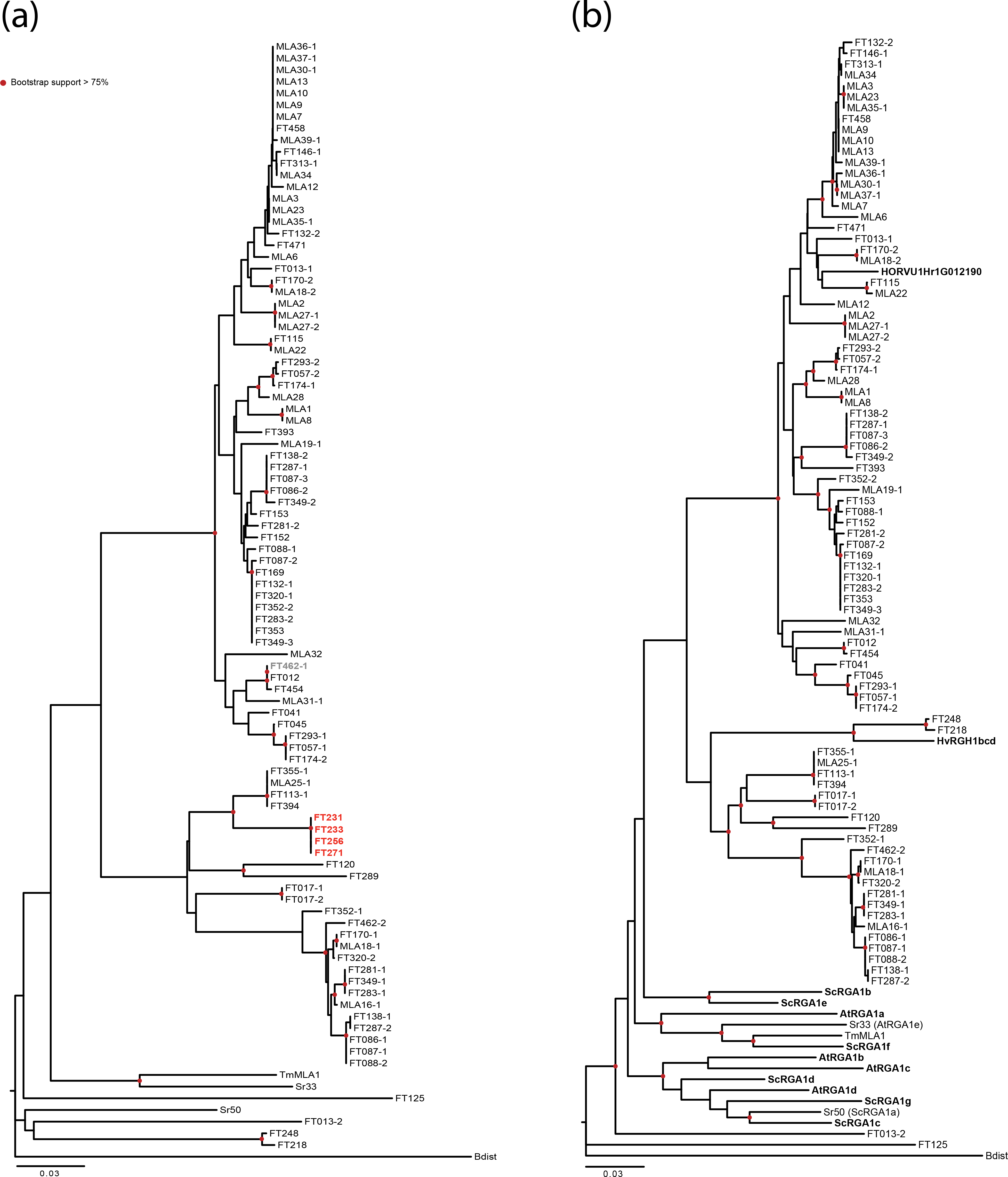
Neighbor-joining (NJ) tree analyses including additional full-length MLA protein sequences. (a) The unrooted NJ tree includes the same 91 amino acid (AA) sequences as shown in fig. 1 plus five additional MLA candidate sequences found in the wild barley accessions. These sequences contain a premature stop at positions 631 (grey) and 561 (red), respectively. (b) The unrooted NJ tree includes the same 91 AA sequences as shown in fig. 1 plus the sequences of two additional MLA homolgs from barley cv. ‘Morex’ (RGH1bcd and a 2nd candidate extracted from the most recent ‘Morex’ assembly) and several additional RGH1 homologs from *Secale cereale* (six sequences) and *Aegilops tauschii* (four sequences). Additional sequences not included in fig. 1 are highlighted in bold. For tree calculation, all positions containing gaps or missing data were omitted (complete deletion option). Red circles mark branches with bootstrap support > 0.75 (500 bootstrap replicates).

**fig. S5:**
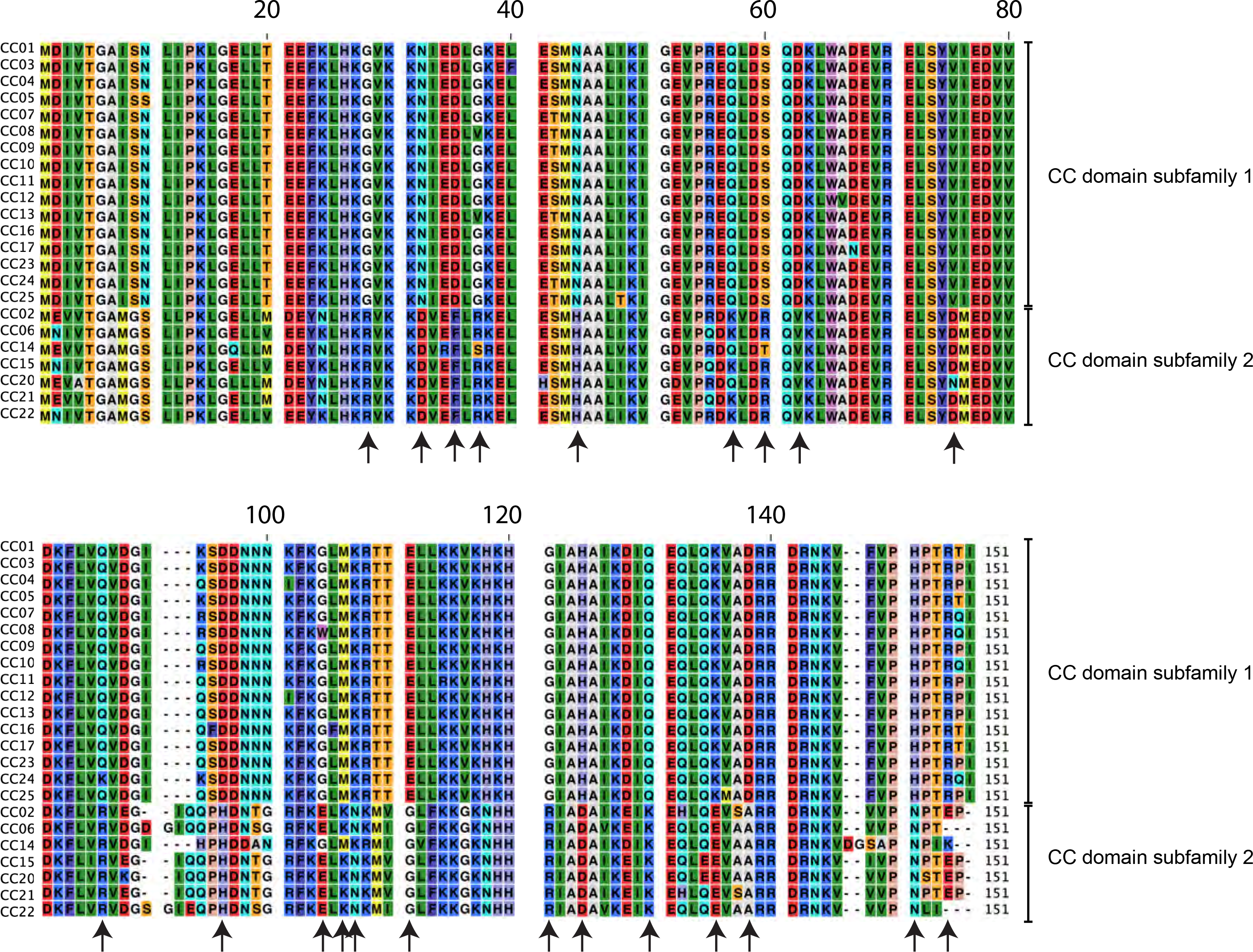
Amino acid sequence alignment of the CC domain haplotypes representing subfamily 1 or subfamily 2 MLA proteins. The alignment shows 22 residues that are differentially occupied by charged or non-charged amino acids in subfamilies 1 and 2 (indicated by arrows).

**fig. S6:**
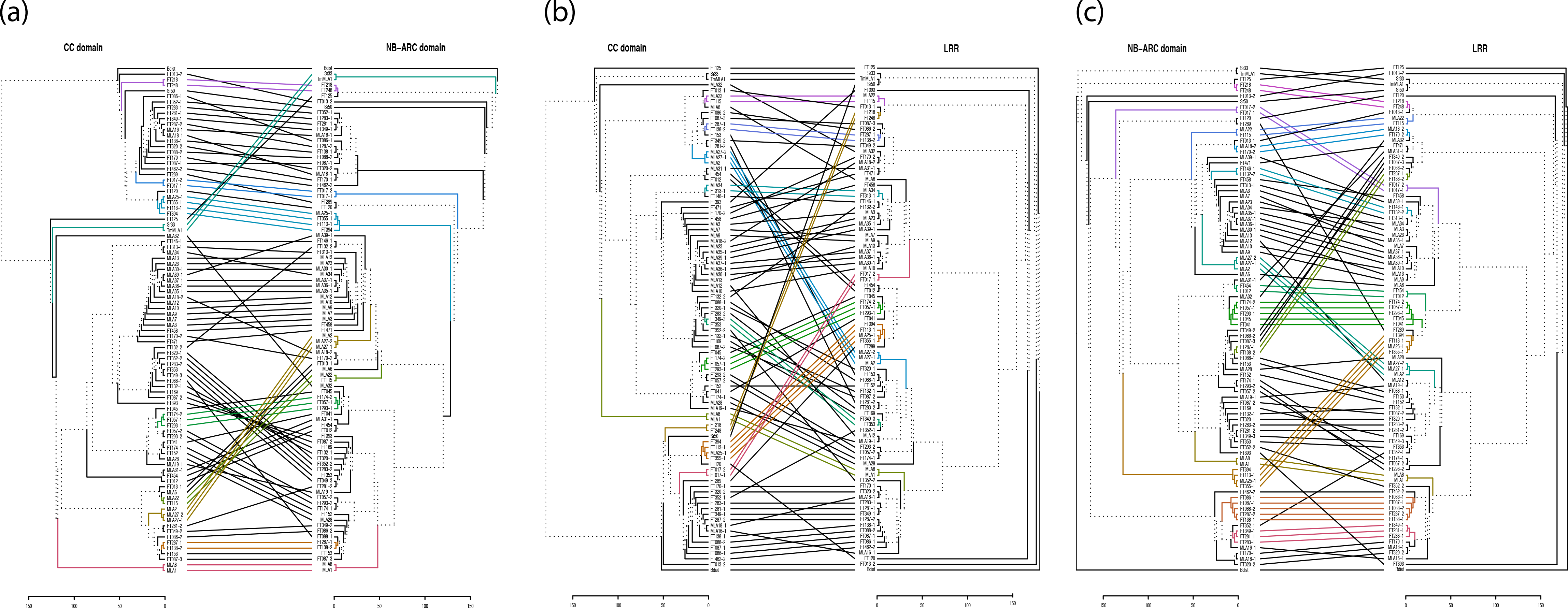
Comparison of neighbor-joining (NJ) trees obtained for the three major MLA protein domains. (a) Comparison of NJ trees obtained for the CC (amino acids 1-151) and NB-ARC (amino acids 180-481) domains. (b) Comparison of NJ trees obtained for the CC domain (amino acids 1-151) and LRR (amino acids 490-end). (c) Comparison of NJ trees obtained for the NB-ARC domain (amino acids 180-481) and LRR (amino acids 490-end). All trees include amino acid sequences of 28 previously published MLAs from barley (Seeholzer et al, 2010), four MLA homologs from other species, and 59 candidate MLA sequences identified in this study from 50 wild barley accessions. All positions containing gaps or missing data were omitted from the tree calculations (complete deletion option). For each comparison, a tanglegram was created from the two corresponding NJ trees using the R package ‘dendextend’, in which lines connect the same isolates and colors represent sub-trees conserved between the two domains. Branches shown as dashed lines indicate differences between the trees.

**fig. S7:**
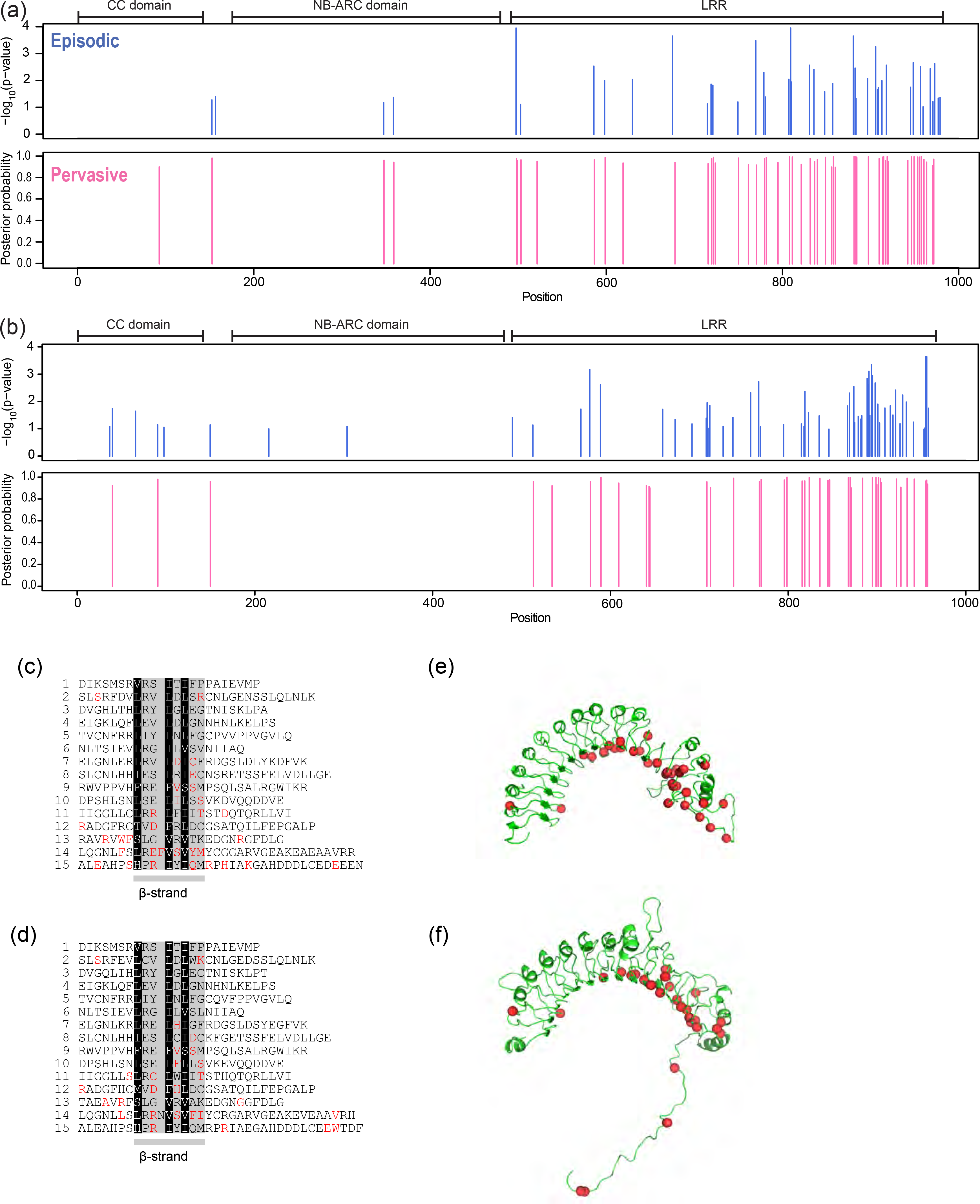
Identification of positively selected sites in MLA conferring resistance to *Bgh* and wild MLA belonging to subfamily 1. (a) Sites under episodic (upper panel; blue bars) or pervasive (lower panel: pink bars) positive selection in a set of 25 known MLA conferring resistance to *Bgh*. (b) Sites under episodic (upper panel; blue bars) or pervasive (lower panel: pink bars) positive selection in 35 newly identified MLAs of wild barley carrying a CC domain belonging to subfamily 1. Only full-length MLA sequences (at least 895 amino acids) from barley were included in these analyses. To test for episodic selection we used MEME and judged all sites with a p value below 0.1 to be under positive selection. To test for pervasive selection, we additionally used FUBAR and judged all sites with a posterior probability above 0.95 to be under positive selection. (c) The deduced secondary structures of the 15 leucine-rich repeat regions (LRR) of MLA1. (d) The deduced secondary structures of the 15 LRR of FT352-2. MLA1 and FT352-2 are selected as representatives of MLA conferring resistance to *Bgh* and the subfamily 1 of wild barley, respectively. The first LRR starts at position 553 for both MLA1 and FT352-2. Amino acid residues under positive selection are indicated by red letters (posterior probability > 0.95). The LxxLxLxx sites, which are proposed to form a short, solvent-exposed β-strand motif (Kajava and Kobe 2002), are indicated. Note that the 14^th^ LRR is irregular with three instead of two x positions after the first L position. Black: hydrophobic core residue; gray: site of any amino acid termed x in the LxxLxLxx motif, (e) The hypothetical tertiary structures of LRR of MLA1 shown in (c) predicted by IntFOLD (McGuffin, et al. 2015). (f) The hypothetical tertiary structures of LRR of FT352-2 shown in (d) predicted by IntFOLD. Amino acid residues under positive selection are indicated by red spheres (posterior probability > 0.95).

**fig. S8:**
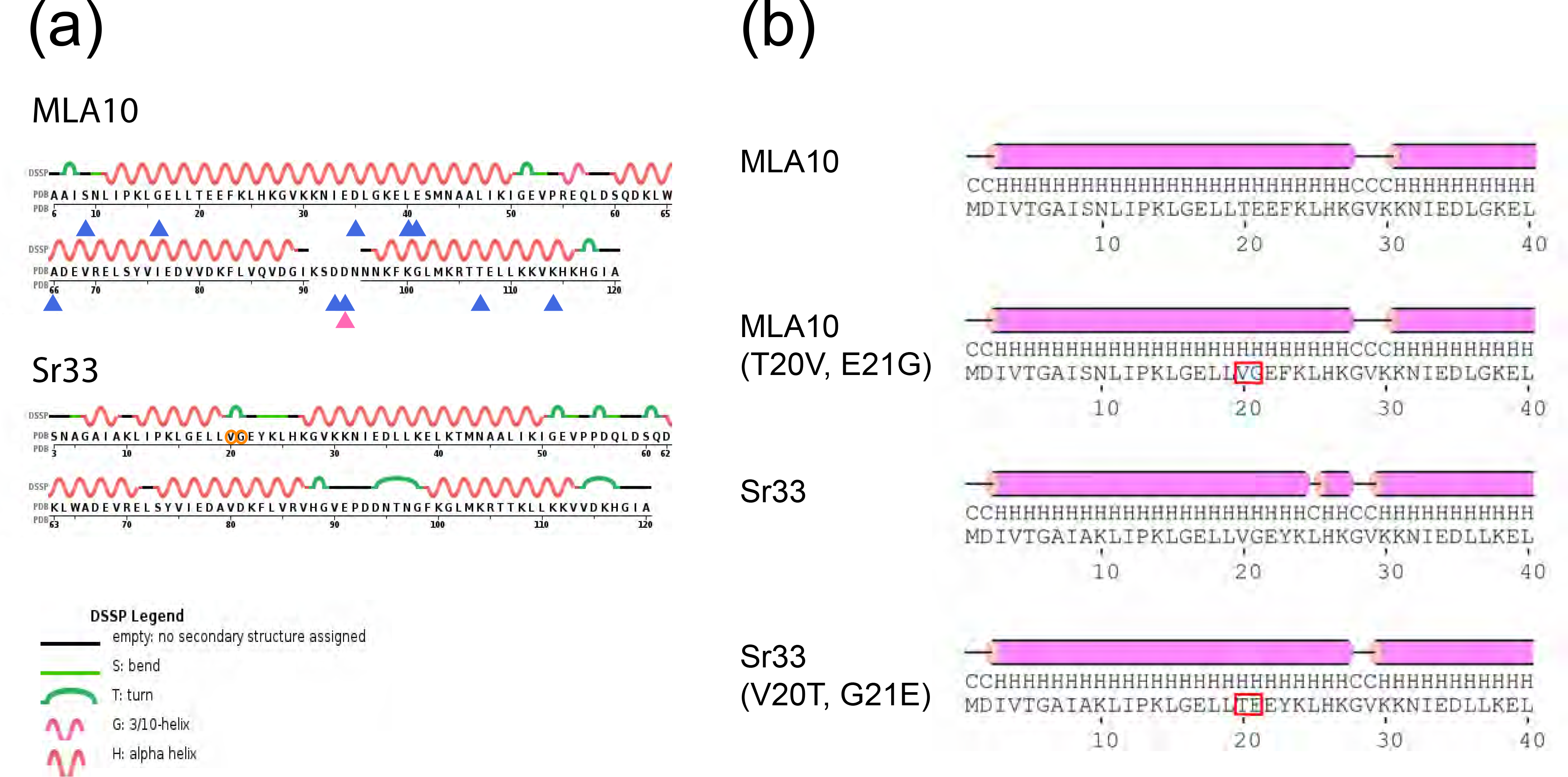
Secondary structure prediction of the mutated coiled-coil (CC) domains of MLA10 and Sr33. (a) Episodically selected sites (blue) and pervasively selected sites (pink) shown in fig. 3A are indicated in the secondary structures of MLA10 (PDB ID 3QFL) and Sr33 (PDB ID 2NCG). The unique residues in the Sr33 CC domain are indicated by orange circles. The schematic secondary structures were obtained from the RCSB Protein Data Bank. (b) Secondary structure prediction for the CC domains (1-40 AA) of MLA10, Sr33, MLA10 (T20V, E21G), and Sr33 (V20T, G20E) using PSIPRED (Buchan DWA, et al. (2013). The red boxes indicate the mutated residues.

### Supplementary tables

**Table S1:** Summary of geographic origin, poulation information and identified MLA sequences for the 50 wild barley lines used in this study

**Table S2:** Number of sequenced read pairs per sample and alignment coverage of identified MLA sequences in 50 wild barley lines

**Table S3:** Summary of unique CC-domain haplotypes (at 100% sequence identity) in wild and domesticated barley

**Table S4:** Summary of nucleotide diversity n for different subsets of MLA and candidate MLA sequences/sequence domains

**Table S5:** The sequences used in the phylogenetic analyses

### Supplementary files

Sequence alignments (fasta) used in figure 1 and 2.

## Materials and Methods

### RNA-Sequencing

Total mRNA from barley plants was obtained from the first or second leaves at 16-19 hours after challenge with *Blumeria graminis* f. sp. *hordei* (*Bgh*) using the RNeasy plant mini kit (Qiagen). RNA-sequencing (RNA-Seq) libraries were prepared by the Max Planck Genome Centre Cologne (Germany) using the Illumina TruSeq stranded RNA sample preparation kit and were subjected to 150-bp paired-end sequencing using the Illumina HiSeq2500 Sequencing System. The RNA-Seq data generated for this study have been deposited in the National Center for Biotechnology Information Sequence Read Archive (SRA) database (BioProject accession no. PRJNA432492, SRA accession no. SRP132475).

### *De novo* transcriptome assembly

For each barley accession, a *de novo* transcriptome assembly was performed using Trinity (version 2.2.0) (Grabherr, et al. 2011) with default parameter settings for paired-end reads; transcript abundance was subsequently estimated (with options --est_method RSEM --algn_method bowtie) and peptide sequences of the best-scoring ORFs were extracted using TransDecoder (Haas, et al. 2013).

### Extraction of MLA candidate sequences

To identify candidate MLA sequences, BLAST searches were performed for the assembled transcripts (from Trinity) and predicted peptides (from TransDecoder) against the CDS and protein sequences of the previously described *Mla* recognition specificities (Seeholzer, et al. 2010) with E value cutoffs of 1e^-6^. For each sample, we identified the Trinity transcript group that contained the best BLAST hit (based on score, identity, and alignment length) against MLA. From this transcript group, a representative candidate MLA transcript was extracted manually based on the BLAST statistics (score, identity, alignment length), transcript abundance (RSEM), inspection of transcript read coverage in the Integrative Genomics Viewer (IGV) (Robinson, et al. 2011), and inspection of a Clustal Omega (Sievers, et al. 2011) multi-sequence alignment (MSA) of the identified and known MLA sequences. Based on this manual inspection, potential sequence errors were corrected according to the RNA-Seq read alignment consensus (IGV visualization) and split sequences were merged where appropriate based on the RNA-Seq read alignment and MSA. Subsequently, the corrected transcripts were used to extract corresponding best-scoring ORFs with TransDecoder, and RNA-Seq reads were mapped against the corrected sequences using bowtie2 (version 2.2.8) (Langmead and Salzberg 2012) with default parameters. The resulting alignments were again inspected in IGV and remaining sequence errors were corrected where required. This procedure of alternating read alignment/ORF prediction and manual screening and error correction was repeated until the RNA-Seq read consensus conformed to the predicted transcript sequence. In the end, from these corrected transcript sequence the final candidate *Mla/MLA* CDS and peptide sequences were extracted with TransDecoder. Accession number for the Mla/MLA sequences from wild barley are listed in Table S5.

### Phylogenetic Analyses

The previously published sequences included in the phylogenetic analyses were retrieved from NCBI (see Table S5 for corresponding accession numbers). As the NCBI accessions for MLA7 (AAQ55540) and MLA10 (AAQ55541) exhibit atypical amino acid variations compared to the previously obtained sequences (Seeholzer, et al. 2010; Maekawa, Cheng, et al. 2011), for these two MLAs we used the corresponding previously published sequences instead of the NCBI entries. The closest MLA homolog in *Brachypodium distachion* was identified by a BLAST search of the MLA consensus sequence against the *Brachypodium* sequences in the non-redundant NCBI protein database. The MLA homologs in the barley cultivar ‘Morex’ were extracted by BLAST searches of the MLA CC-domain against the updated ‘Morex’ genome annotations (Mascher, et al. 2017) (http://webblast.ipk-gatersleben.de/barley_ibsc/).

The MSA visualizations were generated using Unipro UGENE (version) (Okonechnikov, et al. 2012) with Clustal Omega as an alignment tool. The MLA consensus sequence for this visualization was generated from the 25 previously published full-length MLA protein sequences (Seeholzer, et al. 2010). Phylogenetic trees (NJ, ML) were generated using MEGA5 (Tamura, et al. 2011). Neighbor-Net network analysis was performed using SplitsTree4 (applying “proteinMLdist” model: JTT) (Huson and Bryant 2006). CC-domain haplotypes were extracted from an MSA with Clustal Omega. Tanglegrams were generated from the MEGA5 trees (after midpoint rooting) using the ‘tanglegram’ function of the R package ‘dendextend’ (Galili 2015) with previous stepwise greedy rotation to untangle the trees (using function ‘untangle’ with method “step2side”).

For the phylogenetic analyses of individual MLA domains, we regarded the first N-terminal 151 amino acids (1-151) as the CC domain, the sequence stretching from amino acid 180 to 481 as the NB-ARC domain, and the sequence from amino acid 490 to the end as the LRR. The analyses of nucleotide diversity and positive selection were performed on codon alignments of the CDS sequences generated in MEGA5 (using ClustalW as an alignment tool). Nucleotide diversity was then calculated from these alignments using the “nuc.div’ function in the R package ‘pegas’ (Paradis 2010). Sites under episodic positive selection were identified using MEME (Mixed Effects Model of Evolution) (Murrell, et al. 2012) with default parameters and sites under pervasive positive selection were identified using FUBAR (**F**ast, **U**nconstrained **B**ayesian **A**ppRoximation) (Murrell, et al. 2013) with default settings.

### Generation of expression constructs

All CC domains were synthesized by the GeneArt Gene Synthesis service (Thermo Scientific) as pDONR221 entry clones and transferred into the pXCSG-GW-mYFP expression vector (Garcia, et al. 2010) using LR Clonase II (Thermo Scientific). The resulting expression constructs were examined by Sanger sequencing (Eurofins).

### Agrobacterium-mediated transient transformation of *Nicotiana benthamiana* leaves

*Agrobacterium tumefaciens* GV3101 pMP90K were freshly transformed with the respective constructs of interest and grown from single colonies in liquid Luria broth medium containing appropriate antibiotics for ~ 24 hours at 28°C. Bacterial cells were harvested by centrifugation at 2,500 × *g* for 15 min, followed by resuspension in infiltration medium (10 mM MES, pH 5.6, 10 mM MgCl_2_, and 200 μM acetosyringone) to a final OD_600_ = 1.0. Cultures were incubated for 2 to 4 h at 28°C with agitation at 180 rpm before infiltration into leaves of three-to-five-week-old *N. benthamiana* plants. Cell death was assessed three days post infiltration. Experiments were repeated three times independently and the representative image is shown.

### Plant protein extraction and fusion protein detection by immunoblotting

Plant proteins were extracted as described previously (Saur, et al. 2015) with the addition of IGEPAL at a final concentration of 0.25% to the protein extraction buffer. Extracts were diluted 4:1 with 4 x Laemmli buffer (Bio-Rad, 1610747) and heated to 95°C for 5 min. Samples were separated on 10% SDS-PAGE gels, blotted onto PVDF membranes, and probed with anti-GFP (abcam ab6556) followed by anti-rabbit IgG-HRP (Santa Cruz Biotechnology sc-2313) secondary antibodies. Proteins were detected by HRP activity on SuperSignal Femto chemiluminescent substrate (Thermo Fisher 34095) using a Gel Doc™ XR+ Gel Documentation System (Bio-Rad). Experiments were repeated three times independently and the representative image is shown.

### Data Availability Statement

Strains and plasmids are available upon request. Supplemental files are available at FigShare. Sequence data are available at GenBank; the accession numbers are listed in Table S5. The RNA-Seq data generated for this study have been deposited in the National Center for Biotechnology Information Sequence Read Archive (SRA) database (BioProject accession no. PRJNA432492, SRA accession no. SRP132475).

## Acknowledgments

We thank the Max Planck Genome Centre Cologne for RNA-Seq and Petra Köchner and Sabine Haigis for technical assistance. We thank Matthew J. Moscou for sharing unpublished data. We also thank Jijie Chai and Ruben Garrido Oter for helpful suggestions. This work was supported by the Max Planck Society (B.K., I.M.L.S., M.Y.-M., and P.S.-L.) and the German Research Foundation within the scope of the Collaborative Research Centre Grant SFB670 (to B.K., T.M., and P.S.-L.) and German Cluster of Excellence on Plant Sciences (CEPLAS) EXC1028 (to R.K. and M.v.K.). I.M.L.S is supported by a Long-Term Fellowship from The European Molecular Biology Organization (ALTF 368-2016).

## Author contributions

T.M. and P.S.-L. conceived and designed the research; I.M.L.S. and M.Y.-M. performed the research; T.M., B.K., I.M.L.S., and R.K. analyzed data; A.P. and M.v.K. provided materials. T.M., B.K., I.M.L.S., and P.S.-L. wrote the paper with input from all co-authors.

